# Discovery of aryl aminothiazole γ-secretase modulators with novel effects on amyloid β-peptide production

**DOI:** 10.1101/2021.08.09.455471

**Authors:** Sanjay Bhattarai, Lei Liu, Michael S. Wolfe

## Abstract

A series of analogs based on a prototype aryl aminothiazole γ-secretase modulator (GSM) were synthesized and tested for their effects on the profile of 37-to-42-residue amyloid β-peptides (Aβ), generated through processive proteolysis of precursor protein substrate by γ-secretase. Certain substitutions on the terminal aryl D ring resulted in an altered profile of Aβ production compared to that seen with the parent molecule. Small structural changes led to concentration-dependent increases in Aβ37 and Aβ38 production without parallel decreases in their precursors Aβ40 and Aβ42, respectively. The new compounds therefore apparently also stimulate carboxypeptidase trimming of Aβ peptides ≥43 residues, providing novel chemical tools for mechanistic studies of processive proteolysis by γ-secretase.

**Graphical Abstract:** 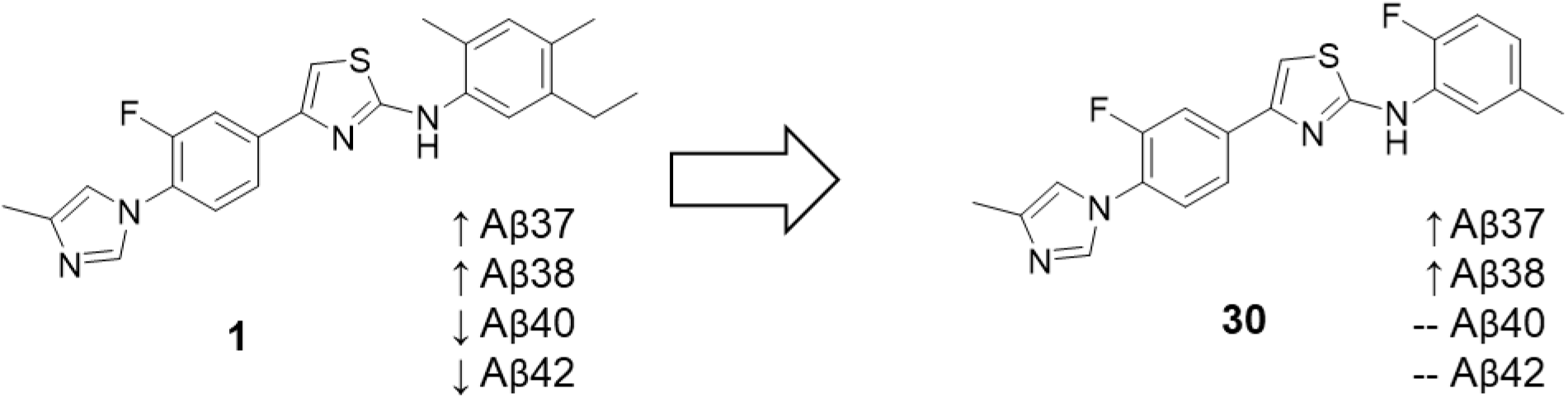

---

Deposition of the 42-residue amyloid β-peptide (Aβ42) as cerebral plaques is a defining pathological feature of Alzheimer’s disease (AD).^1^ Aβ peptides of 37-43 residues are secreted from cells after production through processing of the amyloid precursor protein (APP) by γ-secretase, a membrane-embedded aspartyl protease complex with presenilin as the catalytic component.^2^ More than 200 dominant missense mutations in the genes encoding APP and presenilins are associated with early-onset familial AD.^3^ These mutations, found in the substrate or the enzyme that produces Aβ, alter the production or the properties of Aβ in ways that lead to cerebral plaque deposition. In particular, the ratio of the aggregation-prone Aβ42 to the predominant and more soluble Aβ40 is thought to be the critical factor leading to nucleation into neurotoxic oligomers.^4^ For this reason, the discovery and development of Aβ42-lowering γ-secretase modulators (GSMs) is considered a promising path to therapeutics for the prevention or treatment of AD.^5^

Proteolytic processing of APP substrate within its single transmembrane domain by γ-secretase is complex, involving first endoproteolytic (ε) cleavage to release the APP intracellular domain (AICD) and production of Aβ48 or Aβ49.^6^ These long Aβ peptides, containing most of the APP transmembrane domain are then trimmed—generally in intervals of three amino acids— through a carboxypeptidase function of γ-secretase.^7^ Thus, Aβ production occurs through processive proteolysis along two pathways: Aβ49→Aβ46→Aβ43→Aβ40→Aβ37 and Aβ48→Aβ45→Aβ42→Aβ38. Detailed biochemical analysis of the effects of familial AD mutations in APP and presenilins reveal shifts in ε proteolysis toward proportionally more Aβ48 formation in many cases, resulting in increased Aβ42/Aβ40.^8, 9^ In addition, deficiencies in carboxypeptidase trimming can contribution to increased Aβ42/Aβ40 while also increasing levels of longer Aβ peptide intermediates.^10–12^

Most reported GSMs lower Aβ42 levels and increase Aβ38 levels by stimulating the Aβ42→Aβ38 cleavage event.^5^ Some classes of molecules, including aryl aminothiazoles such as the prototype **1** (**Table 1**), additionally stimulates Aβ40→Aβ37.^13^ The mechanism by which GSMs exert their effects is largely unknown, although one study suggests stabilization of bound Aβ42,^14^ and a new structure of a GSM bound to γ-secretase indicates a binding site on presenilin.^15^ Particularly unclear is whether these compounds selectively stimulate only the final cleavage event or can also affect prior trimming steps. Toward better understanding of processive proteolysis by γ-secretase and the effects of GSMs, we synthesized a series of analogues of **1** to find compounds that might also stimulate other trimming events. Compound **1** was chosen for modification because of its ability to stimulate Aβ40→Aβ37 in addition to Aβ42→Aβ38, demonstrating that stimulation of cleavage events other than Aβ42→Aβ38 is possible.

**Table 1.**
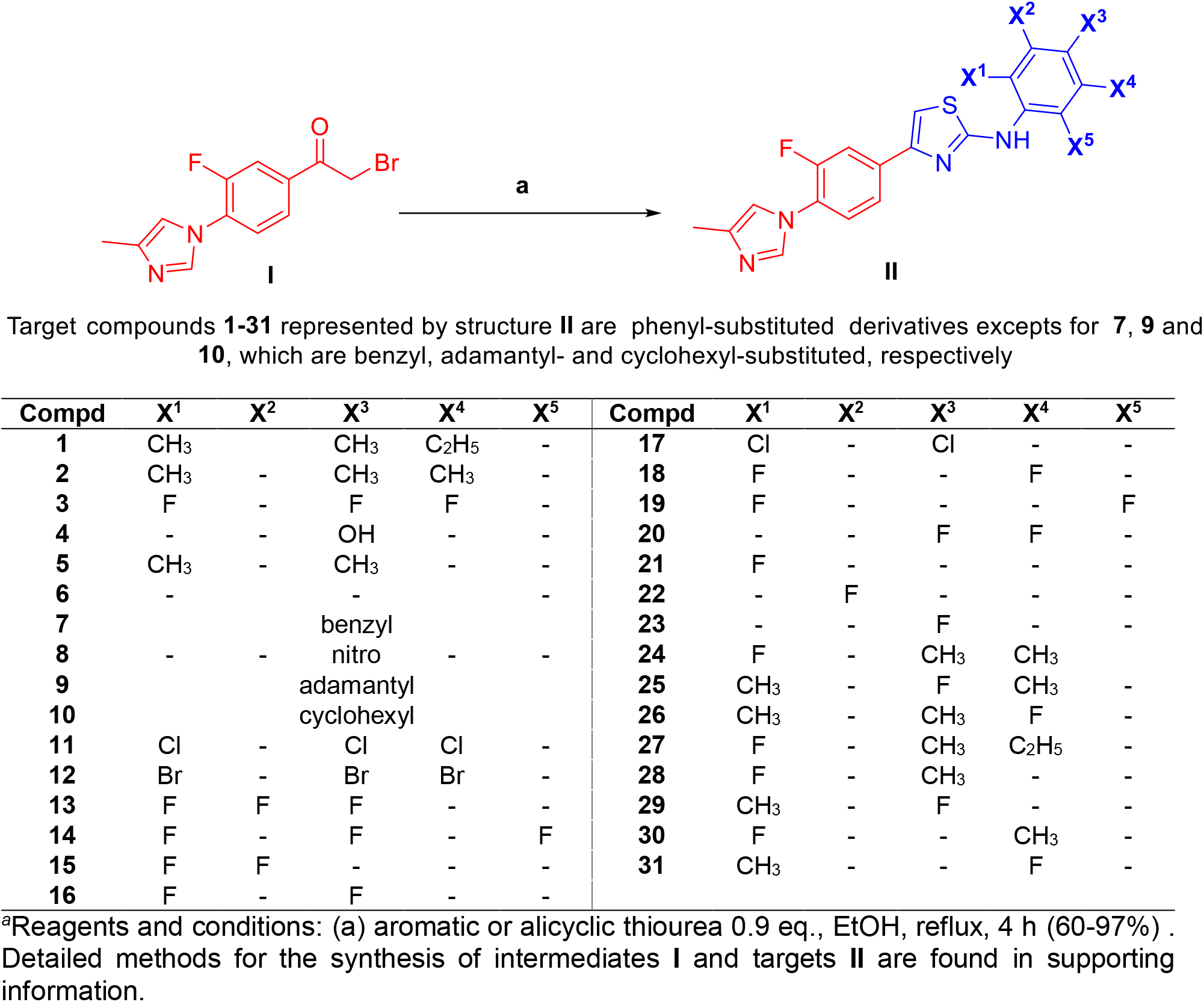
Convergent synthesis of aminothiazole target compounds, D-ring substitution variants of lead compound **1**.

For this initial study attempting to alter the Aβ profile of **1**, we chose the aryl D ring for modification, focusing on removal or replacement of the alkyl groups. Target compounds (**1**-**31**) were synthesized in a convergent synthesis as shown in the scheme in **Table 1** where phenylethanone intermediate **I** was treated with aromatic/alicyclic thioureas under Hantzsch thiazole synthesis condition as previously reported.^16–18^ Briefly for the synthesis of the intermediate **I**, 4-methylimidazole was coupled with 1-(3,4-difluorophenyl)ethan-1-one, and the resulting intermediate was brominated at the keto methyl group to produce **I** in good yields (for details see supporting information). Intermediate aromatic/alicyclic thioureas were synthesized from commercially available nitrates, which were then subjected to Suzuki coupling and/or reduced to amines. The resultant amines were further reacted with benzoyl isothiocyanate to give protected ureas, followed by deprotection of benzoyl group to give intermediate aromatic/alicyclic thioureas in a high yield.

Target compounds 1-31 represented by structure II are phenyl-substituted derivatives excepts for 7, 9 and Compounds **2-10** (**Table 1**) were synthesized first and tested for effects on the profile of secreted peptides Aβ37, Aβ38, Aβ40, and Aβ42. This was performed using human embryonic kidney (HEK) 293 cells transiently transfected with direct γ-secretase substrate C99, the 99-residue C-terminal fragment of APP, with a signal sequence for membrane insertion during translation.^19^ Each of four Aβ peptides secreted into the media was detected using a sensitive and specific two-antibody (“sandwich”) ELISA (**Figure 1**).^19^ The 2,4,5-trimethyl substituted **2** showed a similar effect on the profile of Aβ peptides as the parent compound **1**, only with slightly reduced potency, demonstrating that the 5-ethyl substituent of **1** can be replaced with methyl without much change in potency or profile. However, removal of the 5-methyl of **2** (compound **5**) resulted in >10-fold reduced potency, while removal of all methyl groups (**6**) resulted in dramatically reduced potency. Neither substitution of the phenyl ring in **6** with 4-hydroxyl or 4-nitro substitution (**4**, **8**) nor its replacement with benzyl (**7**), adamantly (**9**), or cyclohexyl (**10**) led to improved activity. Interestingly, replacement of each methyl group in **2** with fluorine (**3**) resulted in an altered effect on the profile of Aβ peptides, increasing Aβ37 levels up to fivefold with little or no effect on Aβ38, Aβ40, or Aβ42.

**Figure 1.**
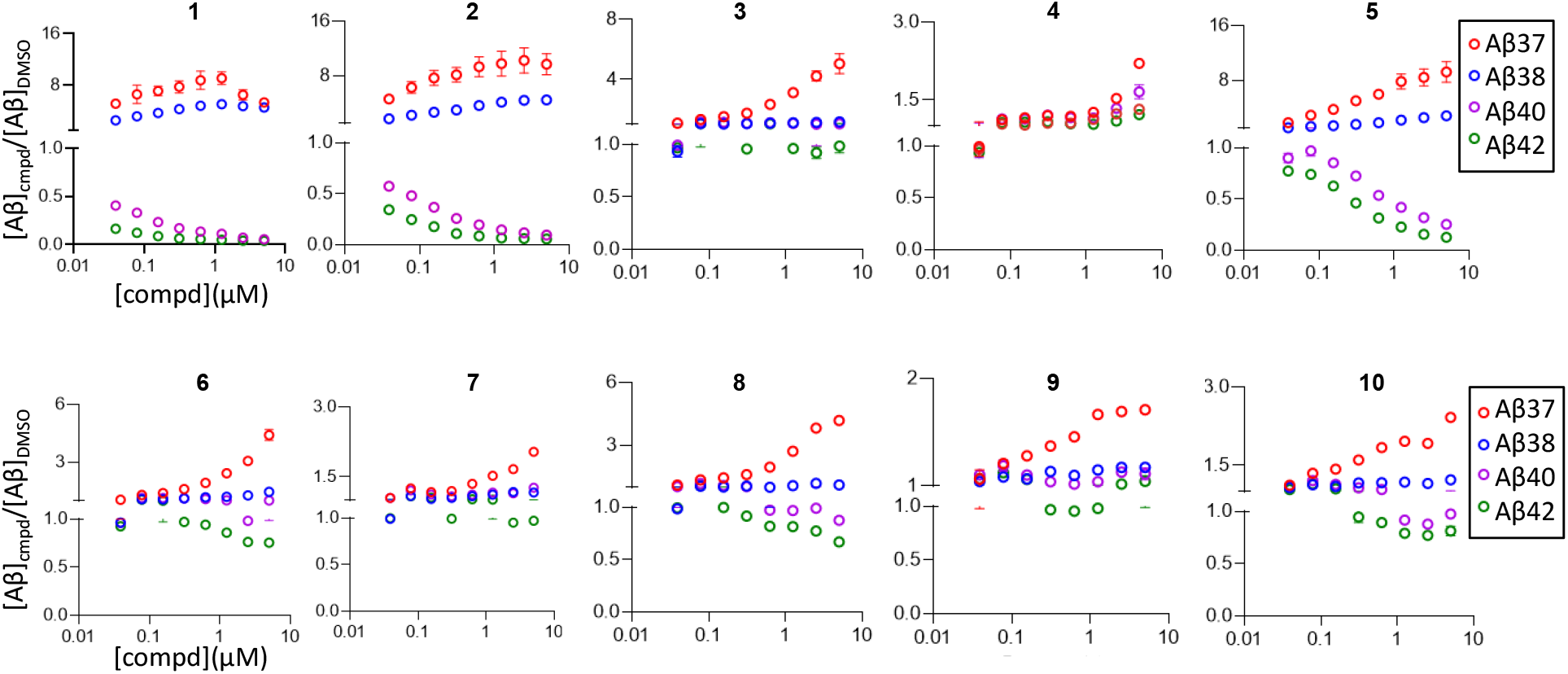
Concentration-dependent effects of **1-10** on Aβ, relative to DMSO vehicle alone, from HEK293 cells transiently transfected with APP C99. Individual Aβ variants (boxed) were measured by specific sandwich ELISAs.

Given the unique effect of **3** on the profile of Aβ peptides, a new series of compounds (**11-31**) was generated to explore the SAR of variations on **3** (**Table 1**). For these analyses, an HEK293 cell line stably expressing C99 with the signal sequence (C99 HEK 293 cells) was generated, to reduce well-to-well variability in Aβ levels associated with transient transfections. Each new compound was tested at a single high concentration (5 μM) for initial evaluation (**Figure 2**). Replacement of the fluorines with chlorine (**11**) resulted in loss of activity, while replacement with bromines (**12**) regained the novel activity profile, increasing Aβ37 and Aβ38 without comparable decreased in Aβ40 and Aβ42. Moving the 5-fluoro of **3** to the 6-position (**14**) led to increased Aβ37 and Aβ38, but now with substantial reduction in Aβ40 and Aβ42, while removal of the 5-fluoro of **3** (compound **16**) increased Aβ37 and Aβ38 as well as Aβ40 and Aβ42. Moving the 4-fluoro group of **16** to the 3-position (**15**) resulted in loss of activity, while replacing the 2- and 4-fluoro groups of **16** with chlorines (**17**) improved the Aβ37 and Aβ38 increases with little or no effects on Aβ40 and Aβ42. Removal of either 2-fluoro or the 4-fluoro of **3** (compounds **20** and **18**, respectively) resulted in loss of activity, and moving the 5-fluoro group of **18** to the 6-position (**19**) did not regain activity. Interestingly though, fluoro substitution alone in either the 2-, 3-, or 4-position (**21**, **22**, or **23**, respectively) led to slightly increased Aβ37 and Aβ38 as well as Aβ40 and Aβ42. Taken together, small differences in the substituent groups and their placement resulted in surprising differences in effects on the profile of Aβ peptides, suggesting that the aryl D ring is tunable in this respect.

**Figure 2.**
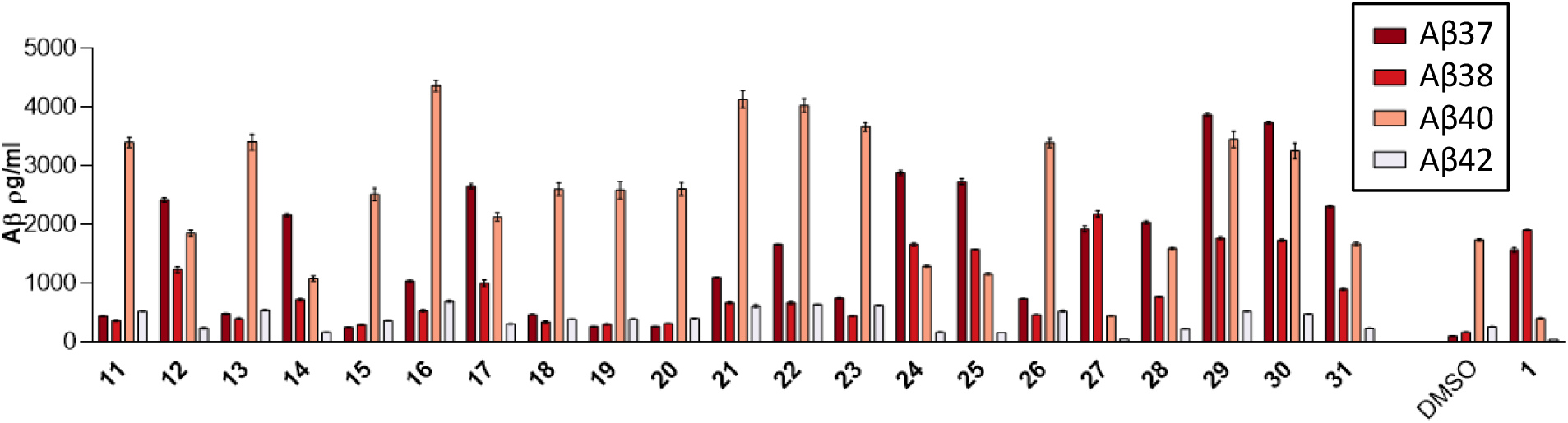
Effects of 5 μM of **11-31** on Aβ from HEK293 cells stably transfected with APP C99, grown in 24-well culture plates. Individual Aβ variants (boxed) were measured by specific sandwich ELISAs. Aβ levels with control DMSO vehicle alone and with 5 μM of parent compound **1** are shown on the right.

A series of analogs were also synthesized to replace only one of the three methyl groups of **2** with fluorine. Replacement of the 2- or 4-methyl group with fluorine (**24** or **25**, respectively) retained the ability to elevate Aβ37 and Aβ38, although with blunted effects on reduction of Aβ40 and Aβ42 compared with **2** (**Figure 2**). In contrast, replacement of the 5-methyl group with fluorine (**26**) resulted in substantial loss of activity. Interestingly, the 5-ethyl counterpart of **24** (compound **27**)—which is also the 2-fluoro counterpart of **1**—displayed stronger effects on lowering Aβ40 and Aβ42 levels compared with **24**. Compounds **28**-**31** represent demethylated counterparts of **24-26**, and each of these compounds retained the ability to increase Aβ37 and Aβ38 without lowering Aβ40 and Aβ42.

Compounds **28**-**31** were selected for dose-response effects in the C99 HEK 293 cell line (**Figure 3**). All compounds showed smooth dose-response effects on elevation of Aβ37 and Aβ38. Each of the new compounds was less effective at elevating Aβ38 compared with prototype **1**. However, compounds **28**-**31** showed stronger effects on Aβ37 elevation compared to **1**, and Aβ37 elevation did not correspond with lowering of precursor Aβ40. This is especially true for **29** and **30**, which showed very little dose-dependent lowering effects on Aβ40. Indeed, as compound concentration increased, Aβ37 levels continued to increase, while Aβ40 levels were lowered slightly and then start to rise again at the maximum 10 μM tested. Similar concentration-dependent effects are seen with Aβ38 and its precursor Aβ42.

**Figure 3.**
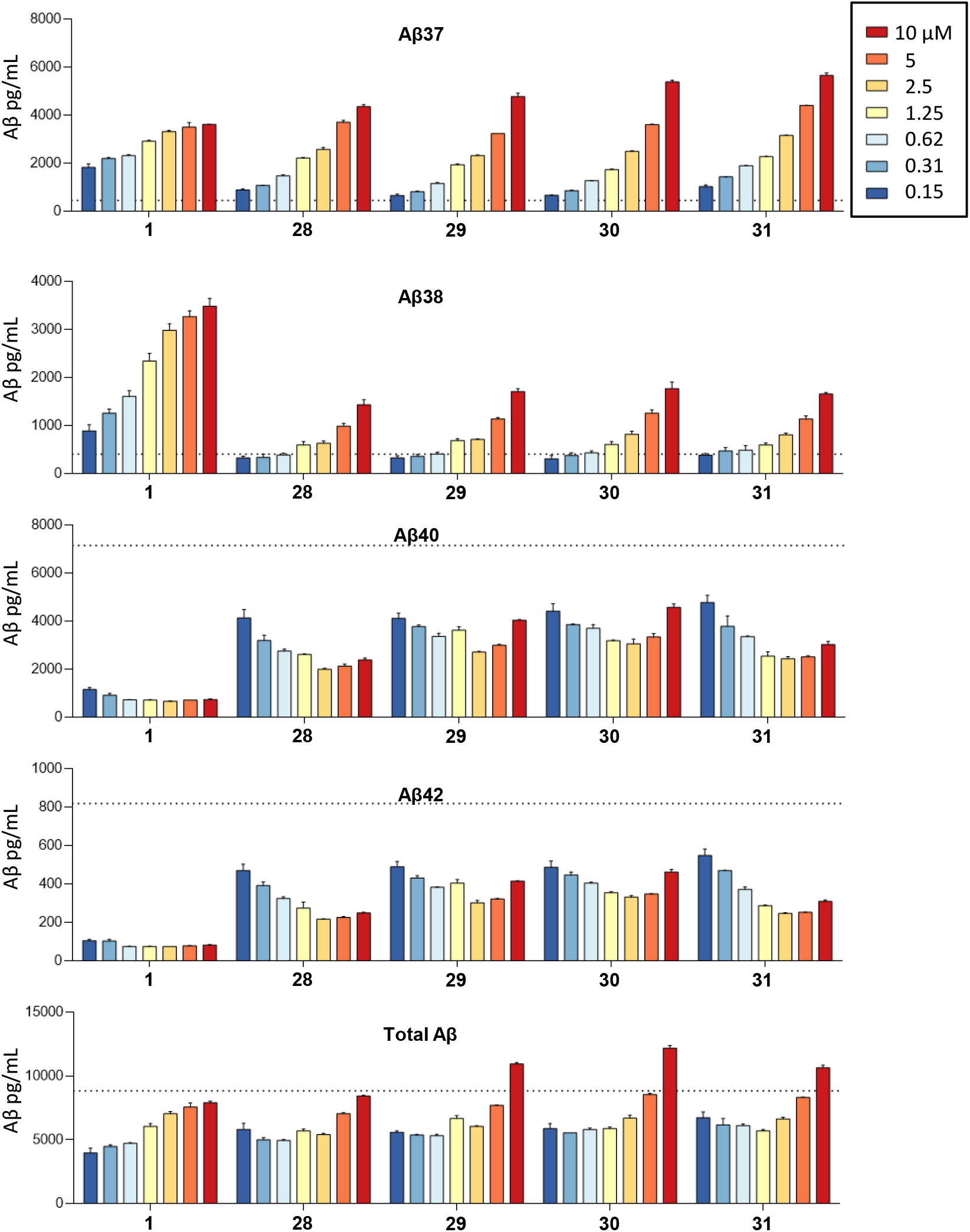
Concentration-dependent effects of **1** and **28-31** on Aβ from HEK293 cells stably transfected with APP C99. Individual Aβ variants (boxed) were measured by specific sandwich ELISAs. Level of each Aβ variant with DMSO vehicle alone is indicated by the dotted lines.

The finding that some of the aryl aminothiazole analogs increase Aβ37 and Aβ38 without comparable lowering of their precursors Aβ40 and Aβ42, respectively, suggests that these compounds are also stimulating prior carboxypeptidase trimming steps along the two processive proteolytic pathways Aβ49→Aβ46→Aβ43→Aβ40→Aβ37 and Aβ48→Aβ45→Aβ42→Aβ38. The compounds do not stimulate overall activity, as levels of endoproteolytic cleavage co-product APP intracellular domain (AICD) are not elevated at the highest concentration test (10 μM), as determined by western blot (data not shown). Total secreted Aβ peptides, detected by ELISA, were not elevated by these compounds relative to DMSO control alone, except for slight elevations with 29, 30, and 31 at the highest concentration (10 μM) (**Figure 3**). Unfortunately, there are no specific ELISAs for the longer Aβ peptide intermediates ranging from Aβ45 to Aβ49. Furthermore, these peptides are membrane-anchored and not normally secreted.^6^ For these reasons, they are difficult to detect in cell-based assays. Specific ELISAs do exist for the secreted Aβ43 peptide;^19^ however, this assay was not sensitive enough for detection in this study, even in our stably transfected C99-expressing cell line (data not shown). Thus, we can only speculate at this point that prior trimming steps are stimulated by the new analogs.

Despite the above limitations, the finding that AICD levels are not elevated by the new analogs implies that total Aβ levels (secreted and intracellular) are also not elevated, as equimolar levels of AICD and total Aβ peptides are produced by γ-secretase.^20^ Therefore, elevations of Aβ37 and Aβ38 should be due to reduction of longer forms of Aβ. For prototype **1**, this is accounted for by reduction of Aβ40 and Aβ42. If lowering of Aβ40 and Aβ42 does not account for increased Aβ37 and Aβ38, this is likely due to stimulation of prior trimming events in addition to Aβ40→Aβ37 and Aβ42→Aβ38. Levels of Aβ40 and Aβ42 represent a balance between their degradation to shorter Aβ peptides and their production from longer Aβ peptides. Stimulation of trimming events involved in both the degradation and production of Aβ40 and Aβ42 would explain their blunted decreases even while end-products Aβ37 and Aβ38 (particularly the former) substantially increase.

We recently reported that 14 different familial AD mutations in the APP transmembrane region are all deficient in one or more carboxypeptidase trimming steps by γ-secretase, leading to elevated Aβ peptides of 45 residues and longer.^11^ These and similar findings with FAD mutations in presenilin^10, 12^ suggest that such longer forms of Aβ may contribute to the pathogenesis of AD.^21^ Compounds that stimulate earlier trimming steps, in addition to the final trimming events Aβ40→Aβ37 Aβ42→Aβ38, might be useful chemical tools to interrogate the pathogenic role of long Aβ peptides and may also have therapeutic potential.

## Supporting information

Supporting Information Available

## Acknowledgment

We thank Dennis Selkoe for helpful discussions. This work was supported by NIH grants GM122894 to M.S.W. and AG15379 to D.J.S.

## Appendix A. Supporting Information.

Supporting Information including detailed chemical and biological methods can be found online.

## References

1. Querfurth HW, LaFerla FM. Alzheimer’s disease. New Eng J Med. 2010;362(4): 329–344.

2. Wolfe MS. Structure and function of the γ-secretase complex. Biochemistry. 2019;58(27): 2953–2966.

3. http://www.alzforum.org/mutations.

4. Selkoe DJ, Hardy J. The amyloid hypothesis of Alzheimer’s disease at 25 years. EMBO Mol Med. 2016;8(6): 595–608.

5. Bursavich MG, Harrison BA, Blain JF. γ-secretase modulators: New Alzheimer’s drugs on the horizon? J Med Chem. 2016;59(16): 7389–7409.

6. Qi-Takahara Y, Morishima-Kawashima M, Tanimura Y, et al. Longer forms of amyloid β protein: implications for the mechanism of intramembrane cleavage by γ-secretase. J Neurosci. 2005;25(2): 436–445.

7. Takami M, Nagashima Y, Sano Y, et al. γ-secretase: Successive tripeptide and tetrapeptide release from the transmembrane domain of β-carboxyl terminal fragment. J Neurosci. 2009;29(41): 13042–13052.

8. Sato T, Dohmae N, Qi Y, et al. Potential link between amyloid β-protein 42 and C-terminal fragment γ49-99 of beta-amyloid precursor protein. J Biol Chem. 2003;278(27): 24294–24301.

9. Bolduc DM, Montagna DR, Seghers MC, Wolfe MS, Selkoe DJ. The amyloid-β forming tripeptide cleavage mechanism of γ-secretase. eLife. 2016;5: pii: e17578.

10. Quintero-Monzon O, Martin MM, Fernandez MA, et al. Dissociation between the processivity and total activity of γ-secretase: Implications for the mechanism of Alzheimer’s disease-causing presenilin mutations. Biochemistry. 2011;50(42): 9023–9035.

11. Devkota S, Williams TD, Wolfe MS. Familial Alzheimer’s disease mutations in amyloid protein precursor alter proteolysis by γ-secretase to increase amyloid β-peptides of >45 residues. J Biol Chem. 2021;296:100281.

12. Fernandez MA, Klutkowski JA, Freret T, Wolfe MS. Alzheimer presenilin-1 mutations dramatically reduce trimming of long amyloid β-peptides (Aβ) by γ-secretase to increase 42-to-40-residue Aβ. J Biol Chem. 2014;289(45): 31043–31052.

13. Kounnas MZ, Danks AM, Cheng S, et al. Modulation of γ-secretase reduces β-amyloid deposition in a transgenic mouse model of Alzheimer’s disease. Neuron. 2010;67(5): 769–780.

14. Szaruga M, Munteanu B, Lismont S, et al. Alzheimer’s-causing mutations shift Aβ length by destabilizing γ-secretase-Aβn interactions. Cell. 2017;170(3): 443–456.

15. Yang G, Zhou R, Guo X, Yan C, Lei J, Shi Y. Structural basis of γ-secretase inhibition and modulation by small molecule drugs. Cell. 2021;184(2): 521–533.

16. Rynearson KD, Buckle RN, Barnes KD, et al. Design and synthesis of aminothiazole modulators of the γ-secretase enzyme. Bioorg Med Chem Lett. 2016;26(16): 3928–3937.

17. Ghodse SM, Telvekar VN. Synthesis of 2-aminothiazole derivatives from easily available thiourea and alkyl/aryl ketones using aqueous NaICl2. Tet Lett. 2015;56(2): 472–474.

18. Bhuniya D, Mukkavilli R, Shivahare R, et al. Aminothiazoles: Hit to lead development to identify antileishmanial agents. Eur J Med Chem. 2015;102:582–93.

19. Liu L, Lauro BM, Wolfe MS, Selkoe DJ. Hydrophilic loop 1 of Presenilin-1 and the APP GxxxG transmembrane motif regulate γ-secretase function in generating Alzheimer-causing Aβ peptides. J Biol Chem. 2021;296:100393.

20. Kakuda N, Funamoto S, Yagishita S, et al. Equimolar production of amyloid β-protein and amyloid precursor protein intracellular domain from β-carboxyl-terminal fragment by γ-secretase. J Biol Chem. 2006;281(21): 14776–14786.

21. Wolfe MS. In search of pathogenic amyloid β-peptide in familial Alzheimer’s disease. Prog Mol Biol Transl Sci. 2019;168:71–78.

