## Supporting Information Available for "Discovery of aryl aminothiazole γ-secretase modulators with novel effects on amyloid β-peptide production"

### SUPPLEMENTAL INFORMATION

### Experimental Section:

#### Biological Protocols

#### Generation and use of HEK293 APP-C99 stably transfected cell line

The cDNA of APP-C99 with N-terminal signal peptide was cloned into pCMV6-AC-IRES-GFP-Puro (OriGene). After transfection of pCMV6-APP-C99-IRES-GFP-Puro into HEK293 cells for 48 h, Puromycin selection was undertaken for 1 week. HEK293 APP-C99 stably transfected cell line were isolated via limiting dilution cloning and confirmed by western blots. This cell line was plated into 24-well plates for initial tests using 5 µM of each compound (shown in Fig. 2) and into 48-well plates for dose-response effects tested for selected compounds (shown in Fig. 3).

#### Tissue culture and transfection of adherent cells

Adherent HEK cells were cultured in complete growth media: Dulbecco's Modified Eagle's Medium (DMEM) supplemented with 10% fetal bovine serum (FBS), 2 mM L-glutamine, 10 units/mL penicillin, and 10 mg/mL streptomycin. For transfection, adherent HEK cells were seeded in 24-well plates at a density of $5x10^5$ cells per well. Transfection was carried out with jetPrime reagent. Cells were incubated for 24 hr and media were changed for conditioning after another 12 hr, at which time the conditioned media were harvested for ELISA, and the cells were harvested for western blots.

#### Aβ ELISA assay

Conditioned media from transfected HEK cells were harvested and diluted with 1% BSA in wash buffer (TBS supplemented with 0.05% Tween). For Aβ 1-x, x-37, x-38, x-40, and x-42 assays, each well of an uncoated 96-well multi-array plate (Meso Scale Discovery, #L15XA-3) was coated with 30 uL of PBS solution containing 3 µg/mL of 266 capture antibody (Elan) and incubated at room temperature overnight. A detection antibody solution was prepared with biotinylated monoclonal antibody recognizing the respective C-terminal residue of each Aβ peptide, plus 100 ng/mL Streptavidin Sulfo-TAG (Meso Scale Discovery, #R32AD-5) and 1% BSA diluted in wash buffer. Following overnight incubation, 50 µL/well of the CM sample plus 25 µL/ well of the detection antibody solution were incubated for 2 h at room temperature with shaking at >300 rpm, washing wells with wash buffer between incubations. The plate was read and analyzed according to manufacturer's protocol.

### General Chemistry

Chemicals and solvents used for organic synthesis were obtained from various commercial vendors (e.g. Fischer, Acros, TCI America, and Sigma-Aldrich). All commercial building blocks and solvents for synthesis were >95% purity and used without further purification or drying. Chemical reactions were monitored via thin layer chromatography (TLC) using aluminum sheets with silica gel 60 $F_{254}$ (Merck). After the completion of reactions, solvents were evaporated by rotavapor and then purified by column chromatography using silica gel 0.060-0.200 mm, pore diameter ca. 6 nm. High-resolution mass spectroscopy (HRMS) analysis was recorded on a LCT Premier mass spectrometer (Micromass Ltd., Manchester, UK), a quadrupole and time-of-flight tandem mass analyzer with an electrospray ion source. The purity analysis of synthesized intermediates as well as final target compounds were measured by LC-MS. For LC/ESI-MS measurement, samples were prepared by dissolving 1 mg/mL of compound in $H_2O$/MeOH (1:1) containing 2 mM ammonium acetate, 10 µL of which was injected for HPLC analysis, eluting with a gradient of water/methanol (containing 2 mM ammonium acetate) from 90:10 to 0:100 for 15 min at a flow rate of 250 µL/min. LC-MS analysis was performed using a Waters Analytical System-Acquity HPLC with an APCI mass spectrometer; UV absorption was detected using an Acquity diode array detector. $^1$H and $^{13}$C NMR spectra were performed on a Bruker Avance 500 and 400 MHz spectrometer and were recorded at ambient temperature using either DMSO-$d_6$, or $CDCl_3$ as solvent. Melting points were measured on Mel-Temp Digital apparatus and are reported without correction.

### Experimental Chemical Methods

**Table S1.** Convergent synthesis of aminothiazole target compounds, D-ring substitution variants of lead compound **1**.

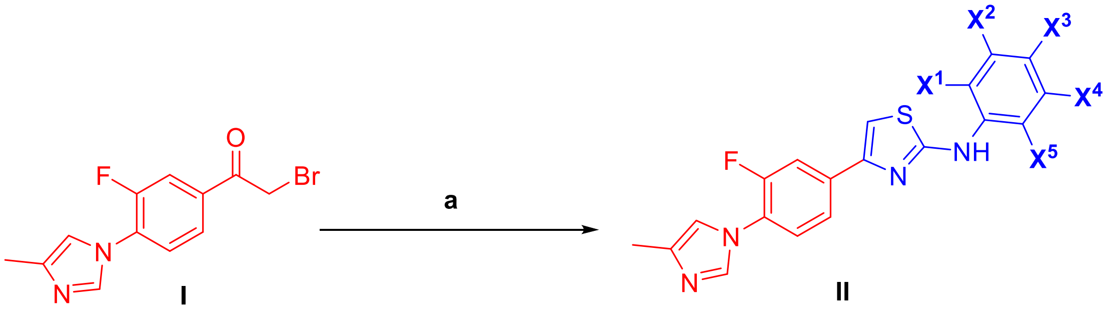


Target compounds **1-31** represented by structure **II** are phenyl-substituted derivatives excepts for **7**, **9** and **10**, which are benzyl, adamantyl- and cyclohexyl-substituted, respectively

| Compd | $X^1$ | $X^2$ | $X^3$ | $X^4$ | $X^5$ | Compd | $X^1$ | $X^2$ | $X^3$ | $X^4$ | $X^5$ |
|---|---|---|---|---|---|---|---|---|---|---|---|
| **1** | $CH_3$ | - | $CH_3$ | $C_2H_5$ | - | **17** | Cl | - | Cl | - | - |
| **2** | $CH_3$ | - | $CH_3$ | $CH_3$ | - | **18** | F | - | - | F | - |
| **3** | F | - | F | F | - | **19** | F | - | - | - | F |
| **4** | - | - | OH | - | - | **20** | - | - | F | F | - |
| **5** | $CH_3$ | - | $CH_3$ | - | - | **21** | F | - | - | - | - |
| **6** | - | | - | | - | **22** | - | F | - | - | - |
| **7** | | | benzyl | | | **23** | - | - | F | - | - |
| **8** | - | - | nitro | - | - | **24** | F | - | $CH_3$ | $CH_3$ | |
| **9** | | | adamantly | | | **25** | $CH_3$ | - | F | $CH_3$ | - |
| **10** | | | cyclohexyl | | | **26** | $CH_3$ | - | $CH_3$ | F | - |
| **11** | Cl | - | Cl | Cl | - | **27** | F | - | $CH_3$ | $C_2H_5$ | - |
| **12** | Br | - | Br | - | Br | **28** | F | - | $CH_3$ | - | - |

| | | | | | | | | | | | |
|---|---|---|---|---|---|---|---|---|---|---|---|
| **13** | F | F | F | - | - | **29** | $CH_3$ | - | F | - | - |
| **14** | F | - | F | - | F | **30** | F | - | - | $CH_3$ | - |
| **15** | F | F | - | - | - | **31** | $CH_3$ | - | - | F | - |
| **16** | F | - | F | - | - | | | | | | |

[a]Reagents and conditions: (a) aromatic or alicyclic thiourea 0.9 eq., EtOH, reflux, 4 h (60-97%). Detailed methods for the synthesis of intermediate **I** and thioureas needed for the synthesis of target compounds (**1**-**31**) are provided below.

#### 2-bromo-1-(3-fluoro-4-(4-methyl-1H-imidazol-1-yl)phenyl)ethan-1-one (I)

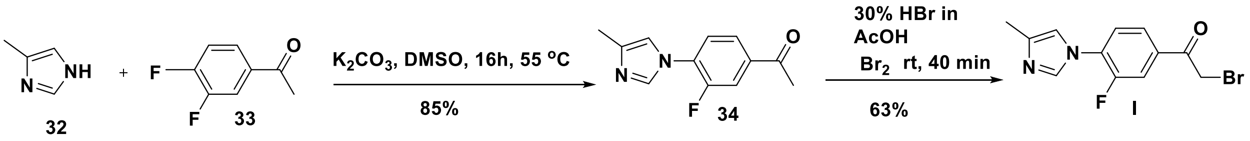


4-Methylimidazole (**32**, 2.0 g, 24.40 mmol) was suspended in DMSO (10 mL), whereupon potassium carbonate (6.74 g, 48.8 mmol) was added. Next, 3,4-Difluoroacetophenone (**33**, 3.38 g, 21.96 mmol) was also added, and the suspension was heated to 55 °C for 16 h. The reaction was cooled to room temperature, then water (10 mL) was added, and the resulting mixture stirred for 1 h at room temperature. Finally, the resulting precipitate collected by filtration was vacuum dried and subjected to column chromatography (ethyl acetate: hexane: 10:90) to afford compound **34** as an orange solid (yield 85%, 4.51 g). To compound **34** (2.0 g, 9.17 mmol) dissolved in ethyl acetate (10 mL), hydrogen bromide (0.8 mL, 33% solution in acetic acid) was added. To this mixture was added bromine [120 µL, 4.5 mmol, in ethyl acetate (10 mL)] drop-wise over 15 minutes at room temperature, and the mixture was stirred for an additional 40 min. After completion of the reaction as indicated by TLC, the reaction mixture was concentrated by rotavapor and subjected to column chromatography (ethyl acetate: hexane: 10:90) provided **I** as an orange-brown solid (1.196 g, 63%).

#### Synthesis of thioureas (35-65)

Thioureas needed for the synthesis of target compounds **1-31** are numbered as **35-65**, respectively**.** The commercially available thoiureas are indicated in the table below with CAS numbers.

**Table S2.** Structures and CAS numbers of commercially available thioureas

| Target compounds | Thioureas | Structure and CAS No. |
|---|---|---|
| **2** | **36** | 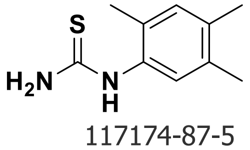 117174-87-5 |
| **3** | **37** | 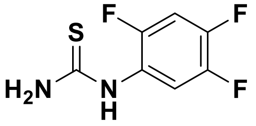 1340519-84-7 |

| | | |
|---|---|---|
| **4** | **38** | 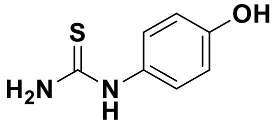 1520-27-0 |
| **5** | **39** | 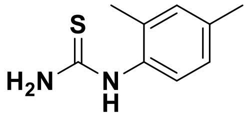 16738-20-8 |
| **6** | **40** | 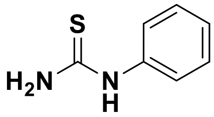 103-85-5 |
| **7** | **41** | 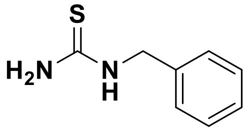 621-83-0 |
| **8** | **42** | 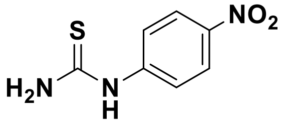 3696-22-8 |
| **9** | **43** | 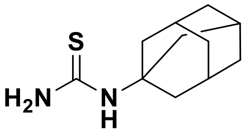 25444-82-0 |
| **10** | **44** | 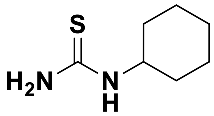 5055-72-1 |
| **11** | **45** | 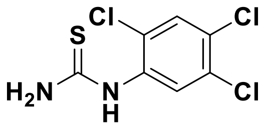 90617-76-8 |
| **12** | **46** | 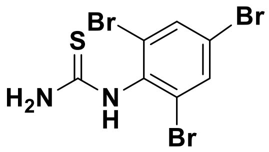 5337-47-3 |
| **13** | **47** | 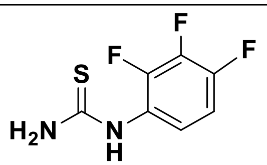 175205-26-2 |

| | | |
|---|---|---|
| **14** | **48** | 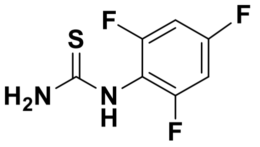 208173-23-3 |
| **15** | **49** | 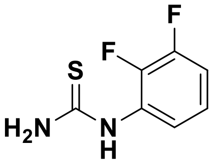 572889-25-9 |
| **16** | **50** | 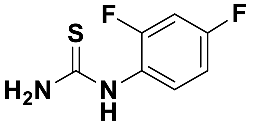 175277-76-6 |
| **17** | **51** | 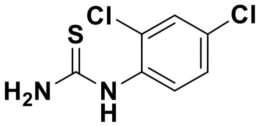 6326-14-3 |
| **18** | **52** | 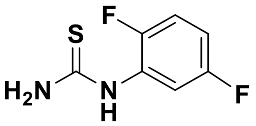 207981-44-0 |
| **19** | **53** | 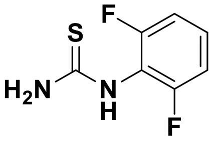 59772-31-5 |
| **20** | **54** | 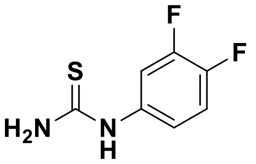 883091-83-6 |
| **21** | **55** | 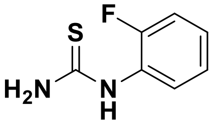 656-32-6 |
| **22** | **56** | 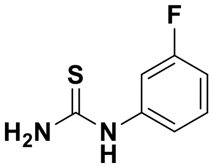 458-05-9 |
| **23** | **57** | 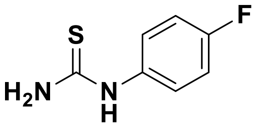 |

| | | |
|---|---|---|
| | | 459-05-2 |
| **28** | **62** | 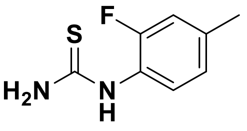 930396-09-1 |
| **29** | **63** | 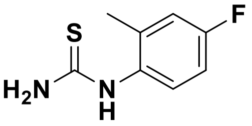 946612-94-8 |
| **30** | **64** | 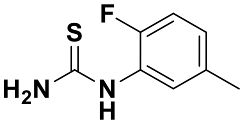 1038356-01-2 |
| **31** | **65** | 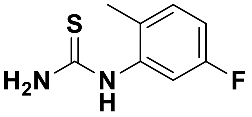 16822-86-9 |

### Synthesis of 1-(5-ethyl-2,4-dimethylphenyl)thiourea (35)

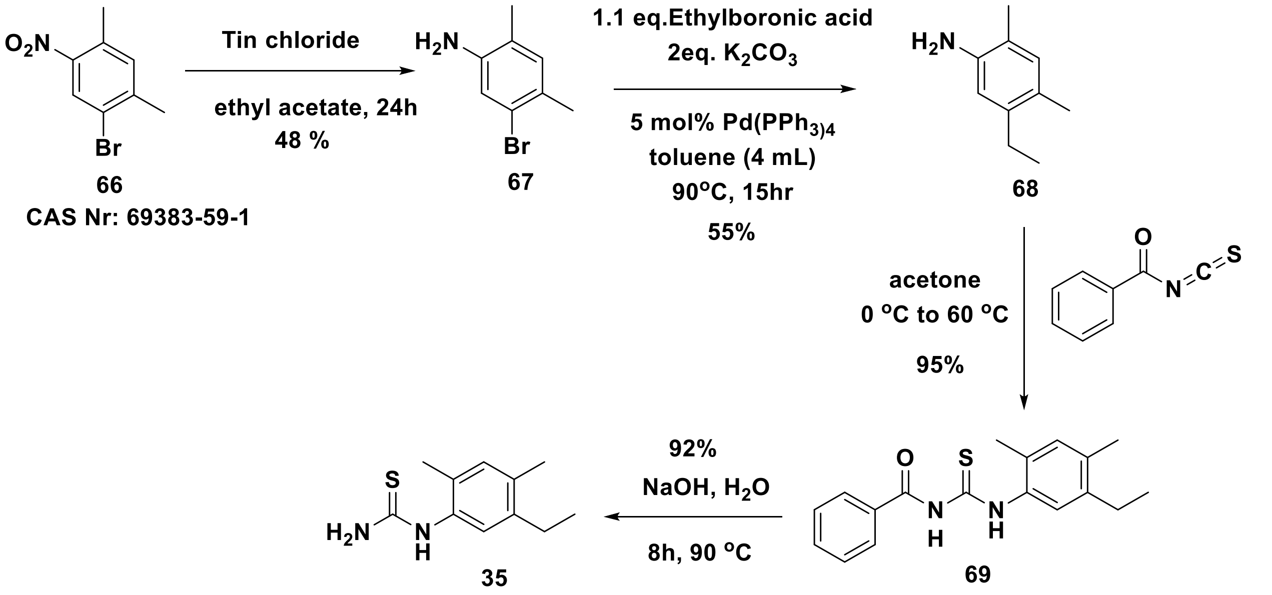

**5-Bromo-2,4-dimethylaniline (67).** 1-Bromo-2,4-dimethyl-5-nitrobenzene (2.0 g, 8.69 mmol) was suspended in ethyl acetate (10 mL). To the resulting suspension, tin (II) chloride (251 mg, 26.08 mmol) was added then further stirred at room temperature at 24 h. After completion of reaction as indicated by TLC, the reaction mixture was concentrated, and then saturated sodium bicarbonate solution (30 mL) was added, followed by extraction of organic layer with ethyl acetate (2 X 50 mL) and washing with brine (2 X 30 mL). The resulting crude material was then subjected to the column chromatography (ethyl acetate: hexane: 5: 95) provided titled compound 5-bromo-2,4-dimethylaniline as an orange-brown solid (834 mg, 48%).

**5-Ethyl-2,4-dimethylaniline (68).** To a solution of 5-bromo-2,4-dimethylaniline (**67**, 800 mg, 3.99 mmol) and ethylboronic acid (443 mg, 6 mmol) in toluene (4 mL) was added $K_2CO_3$ (707.5 mg, 5.12 mmol), and $Pd(PPh_3)_4$ (138 mg, 0.12 mmol). The reaction mixture was heated at 90 °C for 15 h. After completion of reaction as indicated in TLC, the reaction mixture was diluted with $H_2O$ (10 mL) and extracted with EtOAc (2 X 30 mL). The organic layer was separated, dried over anhydrous $Na_2SO_4$ and concentrated by rotavapor. The resultant crude material was purified by column chromatography (ethyl acetate: hexane: 2: 98) to afford **68** (327 mg, 55%) as a white solid.

**N-((5-Ethyl-2,4-dimethylphenyl)carbamothioyl)benzamide** (**69**). To a solution of 5-ethyl-2,4-dimethylaniline (**68**, 300 mg, 2.01 mmol) was added benzoyl isothiocyanate (**70**, 360 mg, 2.21 mmol) and acetone (20 mL) at 0 °C. After 30 min, resulting mixture was heated to 60 °C for 2 h, and the solvent removed under reduced pressure. The resulting white residue was purified by column chromatography (ethyl acetate: hexane: 2: 98) to afford **69** (596 mg, 95%) as a white solid. $^1$H NMR (600 MHz, DMSO-$d_6$) $\delta$ 7.86 – 7.72 (m, 2H), 7.57 – 7.40 (m, 4H), 6.90 – 6.87 (m, 1H), 2.97 – 2.94 (m, 2H), 2.37 (s, 3H), 2.32 (s, 3H), 1.23 – 1.13 (m, 3H). $^{13}$C NMR (150 MHz, DMSO-$d_6$) $\delta$ 179.1, 167.4, 136.7, 135.2, 134.3, 134.1, 128.8, 128.4, 128.3, 125.9, 121.6, 25.7, 18.3, 18.1, 17.5. HRMS (ESI) for $C_{18}H_{20}N_2OSNa$: Calculated: 335.1194, found: 335.1188; Purity by HPLC-UV (214 nm)-ESI-MS: 97.20%. mp 186-188 °C.

**1-(5-Ethyl-2,4-dimethylphenyl)thiourea (35).** N-((5-Ethyl-2,4-dimethylphenyl)carbamothioyl) benzamide (**69**, 500 mg, 1.60 mmol) and sodium hydroxide (128 mg, 3.20 mmol) in water (5 mL) was heated at 90 °C for 8 h, and then concentrated in vacuo. The crude material was dissolved in dichloromethane/water (20 mL each), then washed with brine (30 mL) and extracted with DCM (2 X 50 mL), again concentrated in vacuo and finally purified by column chromatography (ethyl acetate: hexane: 2: 98) to give **35** as a white solid (306 mg, 92%).

**Synthesis of 1-(2-fluoro-4,5-dimethylphenyl)thiourea (58), 1-(4-fluoro-2,5-dimethylphenyl) thiourea (59) and 1-(5-fluoro-2,4-dimethylphenyl)thiourea (60)**

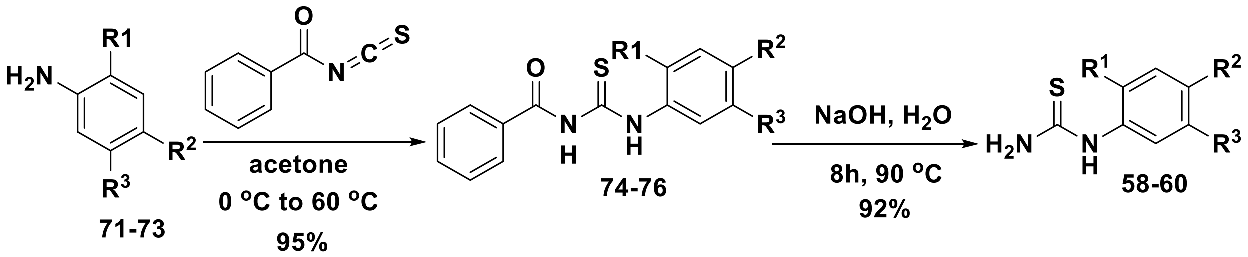


| Compound | $R^1$ | $R^2$ | $R^3$ |
|---|---|---|---|
| **71, 74, 58** | F | $CH_3$ | $CH_3$ |
| **72, 75, 59** | $CH_3$ | F | $CH_3$ |
| **73, 76, 60** | $CH_3$ | $CH_3$ | F |

**General method for the synthesis of phenyl(carbamothioyl)benzamides 74-76**

To a solution of aniline **71**, **72** or **73** (1 eq) was added benzoyl isothiocyanate (**70**, 1.10 eq) and acetone (20 mL) at 0 °C. After 30 min, the mixture was heated to 60 °C for 2 h, and the solvent removed under reduced pressure. The resulting white residue was purified by column chromatography (ethyl acetate: hexane: 2: 98) to afford **74**, **75**, or **76** (95%) as a white solid.

**N-((2-Fluoro-4,5-dimethylphenyl)carbamothioyl)benzamide (74).** Synthesized using 2-fluoro-4,5-dimethylaniline **(71**, 1.0 g, 7.18 mmol) and benzoyl isothiocyanate (**70**, 1.1 g, 7.90 mmol). Yield (2061 mg, 95%). $^{1}$H NMR (600 MHz, DMSO-$d_6$) $\delta$ 7.83 – 7.78 (m, 2H), 7.60 (d, $J$ = 5.1 Hz, 1H), 7.55 – 7.45 (m, 3H), 7.10 (d, $J$ = 8.1 Hz, 1H), 2.32 (s, 3H), 2.31 (s, 3H). $^{13}$C NMR (150 MHz, DMSO-$d_6$) $\delta$ 181.7, 166.3, 154.0, 135.4, 132.3, 129.3, 125.3, 124.9, 120.7, 115.1, 23.5, 21.9, 20.6. HRMS (ESI) for $C_{16}H_{15}FN_2OSNa$: Calculated: 325.0787, found: 325.0825; Purity by HPLC-UV (214 nm)-ESI-MS: 98.50%. mp 178-179 °C.

**N-((4-Fluoro-2,5-dimethylphenyl)carbamothioyl)benzamide (75).** Synthesized using 4-fluoro-2,5-dimethylaniline **(72**, 1.0 g, 7.18 mmol) and benzoyl isothiocyanate (**70**, 1.1 g, 7.90 mmol). Yield (2061 mg, 95%). $^{1}$H NMR (600 MHz, DMSO-$d_6$) $\delta$ 7.82 – 7.77 (m, 2H), 7.61 (d, $J$ = 5.0 Hz, 1H), 7.56 – 7.47 (m, 3H), 7.01 (d, $J$ = 8.0 Hz, 1H), 2.28 (s, 3H), 2.26 (s, 3H). $^{13}$C NMR (150 MHz, DMSO-$d_6$) $\delta$ 182.0, 166.3, 159.7, 134.7, 132.5, 129.1, 127.3, 125.5, 124.0, 122.5, 118.6, 115.2, 23.2, 19.2, 17.0. HRMS (ESI) for $C_{16}H_{15}FN_2OSNa$: Calculated: 325.0787, found: 325.1281; Purity by HPLC-UV (214 nm)-ESI-MS: 96.00%. mp 177-179 °C.

**N-((5-Fluoro-2,4-dimethylphenyl)carbamothioyl)benzamide (76).** Synthesized using 5-fluoro-2,4-dimethylaniline **(73**, 1.0 g, 7.18 mmol) and benzoyl isothiocyanate (**70**, 1.1 g, 7.90 mmol). Yield (2061 mg, 95%). $^{1}$H NMR (600 MHz, DMSO-$d_6$) $\delta$ 7.88 – 7.76 (m, 1H), 7.51 – 7.36 (m, 5H), 7.10 – 7.05 (m, 1H), 2.21 (s, 3H), 2.19 (s, 3H). $^{13}$C NMR (150 MHz, DMSO-$d_6$) $\delta$ 181.0, 166.3, 158.8, 136.5, 135.7, 128.3, 127.7, 124.8, 121.5, , 113.1, 112.7, 21.1, 19.9, 18.3. HRMS (ESI) for $C_{16}H_{15}FN_2OSNa$: Calculated: 325.0787, found: 325.0781; Purity by HPLC-UV (214 nm)-ESI-MS: 97.50%. mp 176-178 °C.

**General method for the synthesis of 1-(2-fluoro-4,5-dimethylphenyl)thiourea (58), 1-(4-fluoro-2,5-dimethylphenyl)thiourea (59) and 1-(5-fluoro-2,4-dimethylphenyl)thiourea (60)**

Phenyl(carbamothioyl)benzamide **74**, **75**, or **76** (1 eq) and sodium hydroxide (2 eq) in water (5 mL) was heated at 90 °C for 8 h, and then concentrated in vacuo. The crude material was dissolved in dichloromethane/water (20 mL each), then washed with brine (30 mL) and extracted with DCM (2 X 50 mL), again concentrated in vacuo and finally purified by column chromatography (ethyl acetate: hexane: 2: 98) to give **58**, **59**, or **60** as a white solid (92%).

**1-(2-Fluoro-4,5-dimethylphenyl)thiourea (58).** Synthesized using N-((2-fluoro-4,5-dimethylphenyl)carbamothioyl)benzamide **(74**, 1.0 g, 3.30 mmol) and sodium hydroxide (264 mg, 6.60 mmol). Yield (603 mg, 92%). $^{1}$H NMR (600 MHz, DMSO-$d_6$) $\delta$ 8.32 (s, 2H), 7.54 (d, $J$ = 5.3 Hz, 1H), 7.03 (d, $J$ = 7.5 Hz, 1H), 3.20 – 3.16 (m, 2H), 2.28 (s, 3H), 2.26 (s, 3H). $^{13}$C NMR (150 MHz, DMSO-$d_6$) $\delta$ 182.5, 153.6, 135.1, 125.9, 121.1, 120.0, 116.9, 21.5, 20.6. HRMS (ESI) for $C_9H_{11}FN_2S$ Na: Calculated: 221.0525, found: 221.0635; Purity by HPLC-UV (214 nm)-ESI-MS: 97.00%. mp 171-172 °C.

**1-(4-Fluoro-2,5-dimethylphenyl)thiourea (59).** Synthesized using N-((4-fluoro-2,5-dimethylphenyl)carbamothioyl)benzamide **(75**, 1.0 g, 3.30 mmol) and sodium hydroxide (264 mg, 6.60 mmol). Yield (603 mg, 92%). $^{1}$H NMR (600 MHz, DMSO-$d_6$) $\delta$ 8.10 (s, 2H), 7.65 (d, $J$ = 5.0 Hz, 1H), 6.98 (d, $J$ = 8.1 Hz, 1H), 2.23 (s, 3H), 2.20 (s, 3H). $^{13}$C NMR (150 MHz, DMSO-$d_6$) $\delta$ 182.1, 159.0, 133.3, 128.9, 125.3, 123.8, 117.2, 23.2, 18.0. HRMS (ESI) for $C_9H_{11}FN_2SNa$:

Calculated: 221.0525, found: 221.0781; Purity by HPLC-UV (214 nm)-ESI-MS: 98.50%. mp 171-172 °C.

**1-(5-Fluoro-2,4-dimethylphenyl)thiourea (60).** Synthesized using N-((5-fluoro-2,4-dimethylphenyl)carbamothioyl)benzamide **(76**, 1.0 g, 3.30 mmol) and sodium hydroxide (264 mg, 6.60 mmol). Yield (603 mg, 92%). $^{1}$H NMR (600 MHz, DMSO-$d_6$) $\delta$ 8.18 (s, 2H), 7.67 (d, $J$ = 8.0 Hz, 1H), 7.13 – 7.09 (m, 1H), 2.28 (s, 3H), 2.19 (s, 3H). $^{13}$C NMR (150 MHz, DMSO-$d_6$) $\delta$ 183.1, 159.6, 135.0, 129.5, 124.9, 120.8, 119.7, 22.1, 19.6. HRMS (ESI) for $C_9H_{11}FN_2SNa$: Calculated: 221.0525, found: 221.1127; Purity by HPLC-UV (214 nm)-ESI-MS: 99.00%. mp 171-172 °C.

#### Synthesis of 1-(5-ethyl-2-fluoro-4-methylphenyl)thiourea (61)

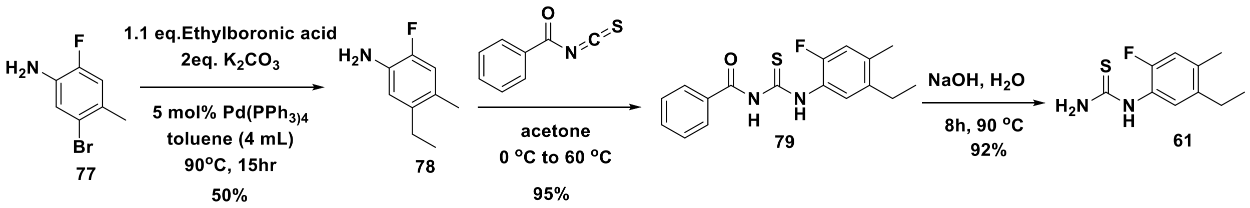


**5-Ethyl-2-flouro-4-dmethylaniline (78).** To the solution of 5-bromo-2-flouro-4-methylaniline **(77**, 1000 mg, 4.90 mmol) and ethylboronic acid (543 mg, 7.35 mmol) in toluene (4 mL) was added $K_2CO_3$ (880 mg, 6.37 mmol), and $Pd(PPh_3)_4$ (283 mg, 0.24 mmol). The reaction mixture was heated at 90 °C for 15h. After completion of reaction as indicated in TLC, the reaction mixture was diluted with $H_2O$ (10 mL) and extracted with EtOAc (2 × 30 mL). The organic layer was separated, dried over anhydrous $Na_2SO_4$ and concentrated in rota vapor. Thus obtained crude was purified by column chromatography (ethyl acetate: hexane: 2: 98) to afford **78** (412 mg, 50%) as a white solid.

**N-((5-Ethyl-2-fluoro-4-methylphenyl)carbamothioyl)benzamide (79).** To a solution of 5-ethyl-2-flouro-4-dmethylaniline (**78**, 400 mg, 2.61 mmol) was added benzoyl isothiocyanate (**70**, 468 mg, 2.87 mmol) and acetone (20 mL) at 0 $^{o}$C. After 30 min, resulting mixture was heated 60 °C for 2 h, and the solvent removed under reduced pressure. The resulting white residue was purified by column chromatography (ethyl acetate: hexane: 2: 98) to afford **79** (784 mg, 95%) as a white solid. $^{1}$H NMR (600 MHz, DMSO-$d_6$) $\delta$ 7.88 – 7.72 (m, 2H), 7.68 (m, 1H), 7.57 – 7.50 (m, 3H), 7.13 (d, $J$ = 7.8 Hz, 1H), 2.53 – 2.36 (m, 2H), 2.21 (s, 3H), 1.13 (t, $J$ = 8.0 Hz, 3H). $^{13}$C NMR (150 MHz, DMSO-$d_6$) $\delta$ 181.3, 166.3, 153.2, 139.5, 134.0, 131.8, 128.1, 126.0, 120.1, 116.9, 20.1, 19.2, 19.0. HRMS (ESI) for $C_{17}H_{17}FN_2OSNa$: Calculated: 339.0944, found: 339.1020; Purity by HPLC-UV (214 nm)-ESI-MS: 99.0%. mp 191-193 °C.

**1-(5-Ethyl-2-fluoro-4-methylphenyl)thiourea (61).** N-((5-Ethyl-2-fluoro-4-methylphenyl) carbamothioyl)benzamide (**79**, 500 mg, 1.58 mmol) and sodium hydroxide (126 mg, 3.16 mmol) in water (5 mL) was heated at 90$^{o}$C for 8 h, and then concentrated in vacuo. The crude material was dissolved in dichloromethane/water (20 mL each), then washed with brine (30 mL) and extracted with DCM (2 X 50 mL), again concentrated in vacuo and finally purified by column chromatography (ethyl acetate: hexane: 2: 98) to give **61** as a white solid (308 mg, 92%). $^{1}$H NMR (600 MHz, DMSO-$d_6$) $\delta$ 8.32 (s, 2H), 7.63 – 7.58 (m, 1H), 7.23 (d, $J$ = 8.1 Hz, 1H), 2.53 (qd, $J$ = 7.9, 1.0 Hz, 2H), 2.21 (s, 3H), 1.13 (t, $J$ = 8.0 Hz, 3H). $^{13}$C NMR (150 MHz, DMSO-$d_6$) $\delta$ 182.5, 152.6, 139.0, 134.5, 126.6, 118.0, 116.6, 20.71, 19.2, 17.6. HRMS (ESI) for $C_{10}H_{13}FN_2SNa$: Calculated: 235.0681, found: 235.0678; Purity by HPLC-UV (214 nm)-ESI-MS: 98.0%. mp 197-199 °C.

**General method for the synthesis of aminothiazole target compounds (1-31).** A solution of 2-bromo-1-(3-fluoro-4-(4-methyl-1*H*-imidazol-1-yl)phenyl)ethanone (**I**, 100 mg, 1 eq) and corresponding thioureas (**35-65**,1 eq) was stirred in absolute ethanol (8 mL) heated to reflux for 4 h. After completion of reaction as indicated in TLC, the reaction mixture was cooled to room temperature, and the solvent removed by rotavapor. The residue was finally purified by column chromatography (DCM: methanol: 2.5: 97.5) to yield **1-31** as solids (60-97%).

**N-(5-Ethyl-2,4-dimethylphenyl)-4-(3-fluoro-4-(4-methyl-1H-imidazol-1-yl)phenyl)thiazol-2-amine (1).** Synthesized using N-((5-fluoro-2,4-dimethylphenyl)carbamothioyl)benzamide **(I**, 100 mg, 0.34 mmol) and 1-(5-ethyl-2,4-dimethylphenyl)thiourea (**35**, 70.80 mg, 0.34 mmol). Yield (125 mg, 92%). $^1$H NMR (600 MHz, DMSO-$d_6$) *δ* 9.37 (s, 1H), 8.01 – 7.71 (m, 3H), 7.70 – 7.56 (m, 2H), 7.43 (s, 1H), 7.30 (q, *J* = 1.0 Hz, 1H), 7.01 (s, 1H), 2.63 – 2.54 (m, 2H), 2.53 – 2.49 (m, 3H), 2.37 (d, *J* = 0.7 Hz, 3H), 2.21 (s, 3H), 2.18 (s, 3H). $^{13}$C NMR (150 MHz, DMSO-$d_6$) δ 166.9, 155.6, 153.6, 148.2, 140.2, 138.1, 136.9, 135.8, 132.6, 131.5, 127.3, 125.7, 124.2, 122.6, 116.5, 114.0, 104.8, 25.7, 18.6, 18.4, 17.8, 15.09, 14.7, 13.9. HRMS (ESI) for $C_{23}H_{23}FN_4SNa$: Calculated: 429.1525, found: 429.1532; Purity by HPLC-UV (214 nm)-ESI-MS: 99%. mp 201-203 °C.

**4-(3-Fluoro-4-(4-methyl-1H-imidazol-1-yl)phenyl)-N-(2,4,5-trimethylphenyl)thiazol-2-amine (2).** Synthesized using N-((5-fluoro-2,4-dimethylphenyl)carbamothioyl)benzamide **(I**, 100 mg, 0.34 mmol) and 1-(2,4,5-trimethylphenyl)thiourea (**36**, 66.05 mg, 0.34 mmol). Yield (129 mg, 98%). $^1$H NMR (600 MHz, DMSO-$d_6$) *δ* 9.39 (s, 1H), 7.88 – 7.59 (m, 3H), 7.57 – 7.45 (m, 2H), 7.33 (s, 1H), 7.22 – 7.16 (m, 1H), 6.97 (s, 1H), 2.43 (s, 3H), 2.35 (d, *J* = 0.7 Hz, 3H), 2.23 (s, 3H), 2.16 (s, 3H). $^{13}$C NMR (150 MHz, DMSO-$d_6$) *δ* 162.5, 155.6, 153.8, 148.4, 140.3, 138.5, 137.9, 136.0, 134.6, 130.9, 129.1, 126.7, 122.1, 116.8,115.2, 110.4, 21.5, 18.1, 17.6, 12.9. HRMS (ESI) for $C_{22}H_{21}FN_4SNa$: Calculated: 415.1369, found: 415.1562; Purity by HPLC-UV (214 nm)-ESI-MS: 99%. mp 203-205 °C.

**4-(3-Fluoro-4-(4-methyl-1H-imidazol-1-yl)phenyl)-N-(2,4,5-trifluorophenyl)thiazol-2-amine (3).** Synthesized using N-((5-fluoro-2,4-dimethylphenyl)carbamothioyl)benzamide **(I**, 100 mg, 0.34 mmol) and 1-(2,4,5-trifluorophenyl)thiourea (**37**, 70.10 mg, 0.34 mmol). Yield (120 mg, 88%). $^1$H NMR (600 MHz, DMSO-$d_6$) *δ* 10.43 (s, 1H), 8.75 – 8.70 (m, 1H), 7.99 – 7.89 (m, 2H), 7.85 (d, *J* = 8.3 Hz, 1H), 7.74 – 7.59 (m, 3H), 7.31 (m, 1H), 2.37 (d, *J* = 0.7 Hz, 3H). $^{13}$C NMR (150 MHz, DMSO-$d_6$) *δ* 162.5, 155.6, 154.1, 153.8, 151.6,147.0, 140.3, 137.9, 130.9, 126.7, 124.7, 122.3, 116.8, 115.2, 110.4, 105.6, 12.9. HRMS (ESI) for $C_{19}H_{12}F_4N_4SNa$: Calculated: 427.0617, found: 427.0781; Purity by HPLC-UV (214 nm)-ESI-MS: 97%. mp 195-196 °C.

**4-(3-Fluoro-4-(4-methyl-1H-imidazol-1-yl)phenyl)-N-(2,4,5-trifluorophenyl)thiazol-2-amine (4).** Synthesized using N-((5-fluoro-2,4-dimethylphenyl)carbamothioyl)benzamide **(I**, 100 mg, 0.34 mmol) and 1-(4-hydroxyphenyl)thiourea (**38**, 57.19 mg, 0.34 mmol). Yield (73.93 mg, 62 %). $^1$H NMR (600 MHz, DMSO-$d_6$) *δ* 9.35 (s, 1H), 8.07 – 8.03 (m, 1H), 7.88 – 7.84 (m, 1H), 7.81 (dd, *J* = 7.5, 5.1 Hz, 1H), 7.59 (dd, *J* = 8.1, 1.4 Hz, 1H), 7.45 (s, 1H), 7.43 – 7.38 (m, 2H), 7.22 – 7.16 (m, 2H), 6.89 – 6.84 (m, 2H), 2.37 (d, *J* = 0.7 Hz, 3H). $^{13}$C NMR (150 MHz, DMSO-$d_6$) *δ* 162.5, 157.5, 155.6, 153.8, 151.6, 151.4, 140.3, 137.9, 135.8, 130.9, 130.3, 126.7, 126.4, 124.7, 124.2, 124.0, 122.3, 122.1, 116.8, 116.5, 115.5, 115.2, 114.4, 110.4, 12.9. HRMS (ESI) for $C_{19}H_{15}FN_4OSNa$: Calculated: 389.0849, found: 389.0912; Purity by HPLC-UV (214 nm)-ESI-MS: 99%. mp 191-192 °C.

**N-(2,4-Dimethylphenyl)-4-(3-fluoro-4-(4-methyl-1H-imidazol-1-yl)phenyl)thiazol-2-amine (5).** Synthesized using N-((5-fluoro-2,4-dimethylphenyl)carbamothioyl)benzamide **(I**, 100 mg, 0.34 mmol) and 1-(2,4-dimethylphenyl)thiourea (**39**, 61.29 mg, 0.34 mmol). Yield (118.70 mg, 95 %). $^1$H NMR (600 MHz, DMSO-$d_6$) *δ* 9.38 (s, 1H), 7.88 – 7.82 (m, 1H), 7.81 (dd, *J* = 7.5, 5.1 Hz, 1H), 7.59 (dd, *J* = 8.1, 1.4 Hz, 1H), 7.45 (s, 1H), 7.24 – 7.19 (m, 2H), 7.19 (dd, *J* = 7.5, 1.4 Hz,

1H), 6.95 – 6.93 (m, 1H), 6.91 (t, *J* = 1.2 Hz, 1H), 2.37 (d, *J* = 0.6 Hz, 3H), 2.21 (s, 3H), 2.16 (s, 3H). $^{13}$C NMR (150 MHz, DMSO-*d*$_6$) *δ* 162.5, 155.6, 153.8, 151.4, 140.3, 137.2, 133.6, 131.8, 128.7, 127.0, 126.4, 124.7, 122.2, 116.8, 115.2, 110.4, 20.7, 19.7, 14.9. HRMS (ESI) for $C_{21}H_{19}FN_4SNa$: Calculated: 401.1212, found: 401.1418; Purity by HPLC-UV (214 nm)-ESI-MS: 99%. mp 203-204 °C.

**4-(3-Fluoro-4-(4-methyl-1H-imidazol-1-yl)phenyl)-N-phenylthiazol-2-amine (6).** Synthesized using N-((5-fluoro-2,4-dimethylphenyl)carbamothioyl)benzamide (**I**, 100 mg, 0.34 mmol) and 1-phenylthiourea (**40**, 51.75 mg, 0.34 mmol). Yield (105 mg, 89 %). $^1$H NMR (600 MHz, DMSO-*d*$_6$) *δ* 9.09 (s, 1H), 7.86 – 7.82 (m, 1H), 7.83 (dd, *J* = 7.5, 5.1 Hz, 1H), 7.69 – 7.62 (m, 2H), 7.58 (dd, *J* = 8.1, 1.5 Hz, 1H), 7.45 (s, 1H), 7.30 – 7.25 (m, 2H), 7.22 – 7.16 (m, 2H), 7.00 (tt, *J* = 7.5, 1.5 Hz, 1H), 2.36 (d, *J* = 0.7 Hz, 3H). $^{13}$C NMR (150 MHz, DMSO-*d*$_6$) *δ* 162.5, 155.6, 153.8, 151.6, 140.3, 139.0, 137.9, 130.9, 129.9, 126.7, 124.7, 123.9, 122.3, 121.6, 116.8, 115.2, 110.4, 12.9. HRMS (ESI) for $C_{19}H_{15}FN_4SNa$: Calculated: 373.0899, found: 373.0913; Purity by HPLC-UV (214 nm)-ESI-MS: 99%. mp 204-205 °C.

**N-Benzyl-4-(3-fluoro-4-(4-methyl-1H-imidazol-1-yl)phenyl)thiazol-2-amine (7).** Synthesized using N-((5-fluoro-2,4-dimethylphenyl)carbamothioyl)benzamide (**I**, 100 mg, 0.34 mmol) and 1-benzylthiourea (**41**, 56.50 mg, 0.34 mmol). Yield (112.70 mg, 92 %). $^1$H NMR (600 MHz, DMSO-*d*$_6$) *δ* 8.29 – 8.20 (m, 1H), 7.94 – 7.92 (m, 1H), 7.85 – 7.81 (m, 1H), 7.78 (dd, *J* = 7.5, 5.1 Hz, 1H), 7.60 (dd, *J* = 8.1, 1.5 Hz, 1H), 7.40 – 7.16 (m, 7H), 4.53 (dt, *J* = 7.5, 1.0 Hz, 2H), 2.18 (d, *J* = 0.6 Hz, 3H). $^{13}$C NMR (150 MHz, DMSO-*d*$_6$) *δ* 166.8, 155.7, 153.6, 148.1, 139.6, 138.0, 136.9, 135.9, 129.0, 128.7, 128.6, 127.8, 127.4, 125.7, 124.0, 122.7, 116.5, 103.8, 48.2, 13.8. HRMS (ESI) for $C_{20}H_{17}FN_4SNa$: Calculated: 387.1056, found: 387.1120; Purity by HPLC-UV (214 nm)-ESI-MS: 99%. mp 206-207 °C.

**4-(3-Fluoro-4-(4-methyl-1H-imidazol-1-yl)phenyl)-N-(4-nitrophenyl)thiazol-2-amine (8). S**ynthesized using N-((5-fluoro-2,4-dimethylphenyl)carbamothioyl)benzamide (**I**, 100 mg, 0.34 mmol) and 1-(4-nitrophenyl)thiourea (**42**, 67 mg, 0.34 mmol). Yield (115.60 mg, 87%). $^1$H NMR (600 MHz, DMSO-*d*$_6$) *δ* 10.29 (s, 1H), 8.33 – 8.25 (m, 1H), 8.22 – 8.16 (m, 1H), 8.07 – 7.98 (m, 1H), 7.89 – 7.82 (m, 4H), 7.78 (s, 1H), 7.69 (dd, *J* = 8.3, 1.4 Hz, 1H), 7.45 – 7.34 (m, 1H), 2.20 (d, *J* = 0.7 Hz, 3H). $^{13}$C NMR (150 MHz, DMSO-*d*$_6$) *δ* 162.5, 155.7, 153.7, 148.7, 147.1, 146.5, 142.6, 140.8, 138.1, 136.9, 135.1, 125.9, 124.6, 123.0, 121.3, 116.6, 114.3, 13.8. HRMS (ESI) for $C_{19}H_{14}FN_5O_2SNa$: Calculated: 418.0750, found: 418.0812; Purity by HPLC-UV (214 nm)-ESI-MS: 98%. mp 203-204 °C.

**4-(3-Fluoro-4-(4-methyl-1H-imidazol-1-yl)phenyl)-N-(4-nitrophenyl)thiazol-2-amine (9).** Synthesized using N-((5-fluoro-2,4-dimethylphenyl)carbamothioyl)benzamide (**I**, 100 mg, 0.34 mmol) and 1-(adamantan-1-yl)thiourea (**43**, 71.40 mg, 0.34 mmol). Yield (89.29 mg, 65%). $^1$H NMR (600 MHz, DMSO-*d*$_6$) *δ* 8.29 – 8.20 (m, 1H), 7.94 – 7.90 (m, 1H), 7.85 (dd, *J* = 7.5, 5.1 Hz, 1H), 7.78 (dd, *J* = 8.1, 1.5 Hz, 1H), 7.60 – 7.54 (m, 1H), 7.34 – 7.31 (m, 5H), 4.53 (s, 1H), 2.18 (d, *J* = 0.6 Hz, 3H), 2.14 – 2.04 (m, 9H), 1.62 (t, *J* = 7.0 Hz, 6H). $^{13}$C NMR (150 MHz, DMSO-*d*$_6$) *δ* 168.8, 155.5, 153.6, 148.1, 139.6, 138.0, 136.9, 135.9, 129.0, 127.8, 127.4, 127.3, 125.7, 124.1, 122.7, 116.5, 114.1, 103.8, 48.2, 42.8, 36.1, 29.2, 13.9. HRMS (ESI) for $C_{23}H_{25}FN_4SNa$: Calculated: 431.1682, found: 431.1732; Purity by HPLC-UV (214 nm)-ESI-MS: 96%. mp 191-192 °C.

**N-Cyclohexyl-4-(3-fluoro-4-(4-methyl-1H-imidazol-1-yl)phenyl)thiazol-2-amine (10).** Synthesized using N-((5-fluoro-2,4-dimethylphenyl)carbamothioyl)benzamide (**I**, 100 mg, 0.34 mmol) and 1-cyclohexylthiourea (**44**, 53.80 mg, 0.34 mmol). Yield (88.70 mg, 74%). $^1$H NMR (600 MHz, DMSO-*d*$_6$) *δ* 7.92 (s, 1H), 7.84 (dd, *J* = 7.5, 5.1 Hz, 1H), 7.77 (dd, *J* = 8.1, 1.5 Hz, 1H), 7.69 (d, *J* = 7.4 Hz, 1H), 7.60 (t, *J* = 8.3 Hz, 1H), 7.28 – 7.16 (m, 1H), 7.25 (s, 1H), 4.90 (d, *J* = 9.7 Hz, 1H), 3.24 – 3.14 (m, 1H), 2.37 (d, *J* = 0.6 Hz, 3H), 1.70 – 1.60 (m, 2H), 1.50 – 1.45 (m, 1H), 1.47

– 1.41 (m, 2H), 1.44 – 1.33 (m, 6H). $^{13}$C NMR (150 MHz, DMSO-$d_6$) $\delta$ 168.0, 155.5, 153.6, 148.2, 138.1, 136.9, 136.1, 136.0, 125.7, 124.0, 123.9, 122.6, 116.5, 114.0, 113.8, 103.0, 53.8, 32.7, 25.8, 24.8, 13.99. HRMS (ESI) for $C_{19}H_{21}FN_4SNa$: Calculated: 379.1369, found: 379.1423; Purity by HPLC-UV (214 nm)-ESI-MS: 99%. mp 199-200 °C.

**4-(3-Fluoro-4-(4-methyl-1H-imidazol-1-yl)phenyl)-N-(2,4,5-trichlorophenyl)thiazol-2-amine (11).** Synthesized using N-((5-fluoro-2,4-dimethylphenyl)carbamothioyl)benzamide (**I**, 100 mg, 0.34 mmol) and 1-(2,4,5-trichlorophenyl)thiourea (**45**, 86.80 mg, 0.34 mmol). Yield (129.80 mg, 85%). $^{1}$H NMR (600 MHz, DMSO-$d_6$) $\delta$ 8.85 (s, 1H), 7.80 (s, 1H), 7.83 (dd, $J$ = 7.5, 5.1 Hz, 1H), 7.60 (s, 1H), 7.65 – 7.58 (m, 2H), 7.55 (s, 1H), 7.28 – 7.18 (m, 2H), 2.39 (d, $J$ = 0.7 Hz, 3H). $^{13}$C NMR (150 MHz, DMSO-$d_6$) $\delta$ 163.5, 156.6, 154.8, 152.6, 152.4, 141.3, 140.5, 138.9, 133.1, 132.0, 131.3, 127.4, 125.7, 123.7, 117.8, 117.5, 116.2, 114.4, 13.9. HRMS (ESI) for $C_{19}H_{12}Cl_3FN_4SNa$: Calculated: 474.9730, found: 475.9830; Purity by HPLC-UV (214 nm)-ESI-MS: 99%. mp 196-197 °C.

**4-(3-Fluoro-4-(4-methyl-1H-imidazol-1-yl)phenyl)-N-(2,4,6-tribromophenyl)thiazol-2-amine (12).** Synthesized using N-((5-fluoro-2,4-dimethylphenyl)carbamothioyl)benzamide (**I**, 100 mg, 0.34 mmol) and 1-(2,4,6-tribromophenyl)thiourea (**46**, 131.16 mg, 0.34 mmol). Yield (126.40 mg, 64%). $^{1}$H NMR (600 MHz, DMSO-$d_6$) $\delta$ 9.90 (s, 1H), 8.09 (s, 2H), 7.92 – 7.85 (m, 1H), 7.79 (dd, $J$ = 8.9, 1.2 Hz, 1H), 7.70 (dd, $J$ = 8.3, 1.9 Hz, 1H), 7.61 (t, $J$ = 8.2, 1.7 Hz, 1H), 7.27 – 7.22 (m, 1H), 2.18 (d, $J$ = 1.0 Hz, 3H). $^{13}$C NMR (150 MHz, DMSO-$d_6$) $\delta$ 166.2, 155.5, 153.5, 148.4, 138.1, 136.9, 135.4, 130.9, 130.3, 125.7, 124.3, 124.2, 122.8, 121.2, 116.4, 113.9, 105.9, 13.9. HRMS (ESI) for $C_{19}H_{12}Br_3FN_4SNa$: Calculated: 606.8215, found: 606.7258; Purity by HPLC-UV (214 nm)-ESI-MS: 98%. mp 193-194 °C.

**4-(3-Fluoro-4-(4-methyl-1H-imidazol-1-yl)phenyl)-N-(2,3,4-triflourophenyl)thiazol-2-amine (13).** Synthesized using N-((5-fluoro-2,4-dimethylphenyl)carbamothioyl)benzamide (**I**, 100 mg, 0.34 mmol) and 1-(2,3,4-trifluorophenyl)thiourea (**47**, 70.10 mg, 0.34 mmol). Yield (120 mg, 88%). $^{1}$H NMR (600 MHz, DMSO-$d_6$) $\delta$ 8.55 (s, 1H), 7.90 (s, 1H), 7.83 (dd, $J$ = 7.5, 5.1 Hz, 1H), 7.57 (dd, $J$ = 8.1, 1.4 Hz, 1H), 7.44 (dt, $J$ = 8.0, 5.0 Hz, 1H), 7.42 (s, 1H), 7.28 – 7.14 (m, 2H), 7.13 (td, $J$ = 8.0, 5.0 Hz, 1H), 2.39 (d, $J$ = 0.7 Hz, 3H). $^{13}$C NMR (150 MHz, DMSO-$d_6$) $\delta$ 163.5, 156.6, 154.8, 148.0, 148.9, 141.3, 138.9, 131.9, 131.3, 126.7, 125.7, 125.2, 123.3, 117.8, 116.5, 113.4, 111.94, 106.6, 14.9. HRMS (ESI) for $C_{19}H_{12}F_4N_4SNa$: Calculated: 427.0617, found: 427.0781; Purity by HPLC-UV (214 nm)-ESI-MS: 97%. mp 195-196 °C.

**4-(3-Fluoro-4-(4-methyl-1H-imidazol-1-yl)phenyl)-N-(2,4,6-trifluorophenyl)thiazol-2-amine (14).** Synthesized using N-((5-fluoro-2,4-dimethylphenyl)carbamothioyl)benzamide (**I**, 100 mg, 0.34 mmol) and 1-(2,4,6-trifluorophenyl)thiourea (**48**, 70.10 mg, 0.34 mmol). Yield (118 mg, 87%). $^{1}$H NMR (600 MHz, DMSO-$d_6$) $\delta$ 8.03 (s, 1H), 7.95 (s, 1H), 7.85 (dd, $J$ = 7.5, 5.1 Hz, 1H), 7.65 (dd, $J$ = 8.1, 1.4 Hz, 1H), 7.55 (s, 1H), 7.32 – 7.26 (m, 2H), 6.75 (td, $J$ = 7.7, 0.9 Hz, 2H), 2.39 (d, $J$ = 0.7 Hz, 3H). $^{13}$C NMR (150 MHz, DMSO-$d_6$) $\delta$ 161.6, 157.4, 153.8, 151.6, 151.4, 140.3, 137.9, 131.3, 127.7, 125.7, 123.3, 117.5, 116.4, 111.4, 105.7, 13.9. HRMS (ESI) for $C_{19}H_{12}F_4N_4SNa$: Calculated: 427.0617, found: 427.0824; Purity by HPLC-UV (214 nm)-ESI-MS: 99%. mp 197-199 °C.

**N-(2,3-Difluorophenyl)-4-(3-fluoro-4-(4-methyl-1H-imidazol-1-yl)phenyl)thiazol-2-amine (15).** Synthesized using N-((5-fluoro-2,4-dimethylphenyl)carbamothioyl)benzamide (**I**, 100 mg, 0.34 mmol) and 1-(2,3-difluorophenyl)thiourea (**49**, 63.90 mg, 0.34 mmol). Yield (109.80 mg, 85%). $^{1}$H NMR (600 MHz, DMSO-$d_6$) $\delta$ 10.38 (s, 1H), 8.55 – 8.26 (m, 1H), 8.02 – 7.94 (m, 2H), 7.94 – 7.80 (m, 2H), 7.76 – 7.54 (m, 1H), 7.40 – 7.19 (m, 2H), 7.13 – 6.89 (m, 1H), 2.19 (d, $J$ = 0.9 Hz, 3H). $^{13}$C NMR (150 MHz, DMSO-$d_6$) $\delta$ 163.4, 155.6, 153.7, 151.4, 149.4, 141.3, 139.3, 131.2, 125.9, 124.5, 122.8, 116.48, 115.3, 114.2, 110.9, 13.9. HRMS (ESI) for $C_{19}H_{13}F_3N_4SNa$:

Calculated: 409.0711, found: 409.0825; Purity by HPLC-UV (214 nm)-ESI-MS: 99%. mp 193-194 °C.

**N-(2,4-Difluorophenyl)-4-(3-fluoro-4-(4-methyl-1H-imidazol-1-yl)phenyl)thiazol-2-amine (16).** Synthesized using N-((5-fluoro-2,4-dimethylphenyl)carbamothioyl)benzamide (**I**, 100 mg, 0.34 mmol) and 1-(2,4-difluorophenyl)thiourea (**50**, 63.90 mg, 0.34 mmol). Yield (109.80 mg, 85%). $^{1}$H NMR (600 MHz, DMSO-$d_6$) $\delta$ 10.14 (s, 1H), 8.54 – 8.50 (m, 1H), 7.99 (d, $J$ = 7.5, 1.7 Hz, 1H), 7.94 (dd, $J$ = 8.1, 1.4 Hz, 1H), 7.86 (dd, $J$ = 8.3, 1.9 Hz, 1H), 7.66 (t, $J$ = 8.3 Hz, 1H), 7.60 (s, 1H), 7.41 – 7.29 (m, 2H), 7.22 – 7.12 (m, 1H), 2.19 (d, $J$ = 1.1 Hz, 3H).$^{13}$C NMR (150 MHz, DMSO-$d_6$) $\delta$ 164.1, 155.6, 153.7, 148.0, 138.0, 136.5, 135.5, 126.0, 125.9, 124.3, 122.8, 121.4, 116.6, 114.2, 111.8, 106.6, 104.3, 13.8. HRMS (ESI) for $C_{19}H_{13}F_3N_4SNa$: Calculated: 409.0711, found: 409.1221; Purity by HPLC-UV (214 nm)-ESI-MS: 98%. mp 194-195 °C.

**N-(2,4-Dichlorophenyl)-4-(3-fluoro-4-(4-methyl-1H-imidazol-1-yl)phenyl)thiazol-2-amine (17).** Synthesized using N-((5-fluoro-2,4-dimethylphenyl)carbamothioyl)benzamide (**I**, 100 mg, 0.34 mmol) and 1-(2,4-dichlorophenyl)thiourea (**51**, 75.10 mg, 0.34 mmol). Yield (98.40 mg, 70%). $^{1}$H NMR (600 MHz, DMSO-$d_6$) $\delta$ 10.70 (s, 1H), 8.16 (d, $J$ = 7.5, 2.3 Hz, 1H), 7.99 – 7.92 (m,2H), 7.87 (dd, $J$ = 8.4, 1.9 Hz, 1H), 7.74 – 7.64 (m, 2H), 7.67 – 7.57 (m, 2H), 7.32 (dd, $J$ = 7.5, 1.5 Hz, 1H), 2.19 (d, $J$ = 1.1 Hz, 3H). $^{13}$C NMR (150 MHz, DMSO-$d_6$) $\delta$ 163.0, 155.6, 153.6, 148.4, 141.3, 138.2, 136.9, 135.2, 131.6, 125.9, 124.6, 118.3, 116.5, 114.1, 106.5, 14.5. HRMS (ESI) for $C_{19}H_{13}Cl_2FN_4SNa$: Calculated: 441.0120, found: 441.1123; Purity by HPLC-UV (214 nm)-ESI-MS: 97%. mp 197-198 °C.

**N-(2,5-Difluorophenyl)-4-(3-fluoro-4-(4-methyl-1H-imidazol-1-yl)phenyl)thiazol-2-amine (18).** Synthesized using N-((5-fluoro-2,4-dimethylphenyl)carbamothioyl)benzamide (**I**, 100 mg, 0.34 mmol) and 1-(2,5-difluorophenyl)thiourea (**52**, 63.90 mg, 0.34 mmol). Yield (109.80 mg, 85%). $^{1}$H NMR (600 MHz, DMSO-$d_6$) $\delta$ 10.43 (s, 1H), 8.56 – 8.50 (m, 1H), 7.99 – 7.87 (m, 2H), 7.86 (dd, $J$ = 8.4, 1.8 Hz, 1H), 7.78 – 7.61 (m, 2H), 7.32 – 7.28 (m, 2H), 6.83 – 6.81 (m, 1H), 2.19 (d, $J$ = 1.1 Hz, 3H). $^{13}$C NMR (150 MHz, DMSO-$d_6$) $\delta$ 163.2, 159.5, 157.6, 155.6, 153.6, 148.9, 147.0, 138.2, 136.9, 135.3, 130.5, 126.0, 124.4, 122.7, 116.5, 114.1, 107.8, 106.1, 13.9. HRMS (ESI) for $C_{19}H_{13}F_3N_4SNa$: Calculated: 409.0711, found: 409.0336; Purity by HPLC-UV (214 nm)-ESI-MS: 99%. mp 194-195 °C.

**N-(2,6-Difluorophenyl)-4-(3-fluoro-4-(4-methyl-1H-imidazol-1-yl)phenyl)thiazol-2-amine (19).** Synthesized using N-((5-fluoro-2,4-dimethylphenyl)carbamothioyl)benzamide (**I**, 100 mg, 0.34 mmol) and 1-(2,6-difluorophenyl)thiourea (**53**, 63.90 mg, 0.34 mmol). Yield (109.80 mg, 85%). $^{1}$H NMR (600 MHz, DMSO-$d_6$) $\delta$ 10.45 (s, 1H), 7.89 – 7.86 (m, 1H), 7.83 (dd, $J$ = 7.5, 5.1 Hz, 1H), 7.58 (dd, $J$ = 8.5, 1.5 Hz, 1H), 7.45 (s, 1H), 7.22 – 7.16 (m, 2H), 7.11 – 7.05 (m, 1H), 6.98 – 6.95 (m, 2H), 2.37 (d, $J$ = 1.0 Hz, 3H). $^{13}$C NMR (150 MHz, DMSO-$d_6$) $\delta$ 162.6, 158.9, 155.6, 154.8, 151.6, 140.3, 137.9, 131.9, 130.3, 129.5, 125.7, 124.4, 123.3, 121.1, 116.8, 115.2, 114.4, 110.4, 12.89. HRMS (ESI) for $C_{19}H_{13}F_3N_4SNa$: Calculated: 409.0711, found: 409.0825; Purity by HPLC-UV (214 nm)-ESI-MS: 98%. mp 194-195 °C.

**N-(3,4-Difluorophenyl)-4-(3-fluoro-4-(4-methyl-1H-imidazol-1-yl)phenyl)thiazol-2-amine (20).** Synthesized using N-((5-fluoro-2,4-dimethylphenyl)carbamothioyl)benzamide (**I**, 100 mg, 0.34 mmol) and 1-(3, 4-difluorophenyl)thiourea (**54**, 63.90 mg, 0.34 mmol). Yield (109.80 mg, 85%). $^{1}$H NMR (600 MHz, DMSO-$d_6$) $\delta$ 9.54 (s, 1H), 7.88 – 7.81 (m, 1H), 7.81 (dd, $J$ = 7.5, 5.1 Hz, 1H), 7.59 (dd, $J$ = 8.1, 1.4 Hz, 1H), 7.45 (s, 1H), 7.38 – 7.34 (m, 1H), 7.32 – 7.26 (m, 1H), 7.22 – 7.16 (m, 3H), 2.39 (d, $J$ = 0.7 Hz, 3H). $^{13}$C NMR (150 MHz, DMSO-$d_6$) δ 162.5, 155.6, 153.8, 151.6, 149.1, 140.3, 137.9, 135.6, 130.9, 126.7, 124.7, 122.3, 119.9, 116.0, 114.4, 110.4, 107.8, 13.9. HRMS (ESI) for $C_{19}H_{13}F_3N_4SNa$: Calculated: 409.0711, found: 409.0721; Purity by HPLC-UV (214 nm)-ESI-MS: 98%. mp 192-193 °C.

**N-(2-Fluorophenyl)-4-(3-fluoro-4-(4-methyl-1H-imidazol-1-yl)phenyl)thiazol-2-amine (21).** Synthesized using N-((5-fluoro-2,4-dimethylphenyl)carbamothioyl)benzamide (**I**, 100 mg, 0.34 mmol) and 1-(2-fluorophenyl)thiourea (**55**, 57.80 mg, 0.34 mmol). Yield (104.80 mg, 84%). $^{1}$H NMR (600 MHz, DMSO-$d_6$) $\delta$ 10.17 (s, 1H), 8.59 – 8.57 (m, 1H), 7.99 – 7.91 (m, 2H), 7.69 – 7.65 (m, 1H), 6.61 (s, 1H), 7.34 – 7.22 (m, 3H), 7.09 – 6.99 (m, 1H), 2.19 (d, *J* = 1.0 Hz, 3H). $^{13}$C NMR (150 MHz, DMSO-$d_6$) $\delta$ 163.9, 155.6, 153.7, 151.1, 148.1, 138.2, 136.9, 135.5, 126.4, 125.9, 124.4, 122.8, 122.0, 116.48, 116.5, 115.6, 114.2, 106.7, 14.5. HRMS (ESI) for $C_{19}H_{14}F_2N_4SNa$: Calculated: 391.0805, found: 391.0921; Purity by HPLC-UV (214 nm)-ESI-MS: 99%. mp 198-199 °C.

**N-(3-Fluorophenyl)-4-(3-fluoro-4-(4-methyl-1H-imidazol-1-yl)phenyl)thiazol-2-amine (22).** Synthesized using N-((5-fluoro-2,4-dimethylphenyl)carbamothioyl)benzamide (**I**, 100 mg, 0.34 mmol) and 1-(3-fluorophenyl)thiourea (**56**, 57.80 mg, 0.34 mmol). Yield (104.80 mg, 84%). $^{1}$H NMR (600 MHz, DMSO-$d_6$) $\delta$ 10.61 (s, 1H), 8.00 – 7.91 (m, 2H), 7.88 (dd, *J* = 8.4, 1.9 Hz, 1H), 7.84 – 7.75 (m, 1H), 7.73 – 7.68 (m, 1H), 7.64 (s, 1H), 7.41 – 7.36 (m, 2H), 7.35 – 7.28 (m, 1H), 6.82 – 6.78 (m, 1H), 2.19 (d, *J* = 1.1 Hz, 3H). $^{13}$C NMR (150 MHz, DMSO-$d_6$) $\delta$ 164.0, 162.0, 155.6, 153.7, 148.4, 143.1, 138.2, 136.9, 135.4, 131.0, 125.9, 124.4, 122.8, 116.5, 114.2, 113.3, 108.1, 106.1, 13.9. HRMS (ESI) for $C_{19}H_{14}F_2N_4SNa$: Calculated: 391.0805, found: 391.0854; Purity by HPLC-UV (214 nm)-ESI-MS: 99%. mp 197-196 °C.

**N-(4-Fluorophenyl)-4-(3-fluoro-4-(4-methyl-1H-imidazol-1-yl)phenyl)thiazol-2-amine (23).** Synthesized using N-((5-fluoro-2,4-dimethylphenyl)carbamothioyl)benzamide (**I**, 100 mg, 0.34 mmol) and 1-(4-fluorophenyl)thiourea (**57**, 57.80 mg, 0.34 mmol). Yield (104.80 mg, 84%). $^{1}$H NMR (600 MHz, DMSO-$d_6$) $\delta$ 10.38 (s, 1H), 8,00 – 7.92 (m, 2H), 7.88 (dd, *J* = 8.3, 1.9 Hz, 1H), 7.82 – 7.71 (m, 1H), 7.69 – 7.67 (m, 1H), 7.56 (s, 1H), 7.33 – 7.29 (m, 1H), 7.24 – 7.14 (m, 2H), 2.19 (d, *J* = 1.0 Hz, 3H). $^{13}$C NMR (150 MHz, DMSO-$d_6$) $\delta$ 163.8, 158.3, 156.4, 155.6, 153.7, 148.3, 138.1, 136.9, 135.5, 125.8, 124.4, 122.8, 118.9, 116.9, 115.8, 114.2, 105.5, 14.5. HRMS (ESI) for $C_{19}H_{14}F_2N_4SNa$: Calculated: 391.0805, found: 391.0264; Purity by HPLC-UV (214 nm)-ESI-MS: 99%. mp 196-197 °C.

**N-(2-Fluoro-4,5-dimethylphenyl)-4-(3-fluoro-4-(4-methyl-1H-imidazol-1-yl)phenyl)thiazol-2-amine (24).** Synthesized using N-((5-fluoro-2,4-dimethylphenyl)carbamothioyl)benzamide (**I**, 100 mg, 0.34 mmol) and 1-(2-fluoro-4,5-dimethylphenyl)thiourea (**58**, 67.40 mg, 0.34 mmol). Yield (104.80 mg, 84%). $^{1}$H NMR (600 MHz, DMSO-$d_6$) $\delta$ 9.93 (s, 1H), 8.16 (d, *J* = 8.6 Hz, 1H), 7.97 – 7.93 (m, 1H), 7.91 (dd, *J* = 12.6, 1.9 Hz, 1H), 7.84 (dd, *J* = 8.3, 1.9 Hz, 1H), 7.69 – 7.64 (m, 1H), 7.53 (s, 1H), 7.32 – 7.28 (m, 1H), 7.07 (d, *J* = 12.1 Hz, 1H), 2.25 (s, 3H), 2.23 – 2.14 (m, 6H). $^{13}$C NMR (150 MHz, DMSO-$d_6$) $\delta$ 164.6, 155.6, 153.6, 151.9, 149.9, 138.1, 136.9, 132.5, 131.7, 126.4, 125.8, 124.3, 122.7, 116.6, 114.1, 113.9, 106.0, 19.7, 19.2, 13.9. HRMS (ESI) for $C_{21}H_{18}F_2N_4SNa$: Calculated: 392.1118, found: 392.1231; Purity by HPLC-UV (214 nm)-ESI-MS: 98%. mp 207-208 °C.

**N-(4-Fluoro-2,5-dimethylphenyl)-4-(3-fluoro-4-(4-methyl-1H-imidazol-1-yl)phenyl)thiazol-2-amine (25).** Synthesized using N-((5-fluoro-2,4-dimethylphenyl)carbamothioyl)benzamide (**I**, 100 mg, 0.34 mmol) and 1-(4-fluoro-2,5-dimethylphenyl)thiourea (**59**, 67.40 mg, 0.34 mmol). Yield (104.80 mg, 84%). $^{1}$H NMR (600 MHz, DMSO-$d_6$) $\delta$ 9.95 (s, 1H), 8.14 (dd, *J* = 7.5, 5.1 Hz, 1H), 7.95 (dd, *J* = 8.1, 1.4 Hz, 1H), 7.85 (s, 1H), 7.81 (d, *J* = 4.8 Hz, 1H), 7.62 – 7.56 (m, 2H), 7.32 – 6.05 (m, 2H), 2.37 (d, *J* = 0.7 Hz, 3H), 2.26 (s, 3H), 2.22 (s, 3H). $^{13}$C NMR (150 MHz, DMSO-$d_6$) $\delta$ 162.5, 159.0, 155.6, 153.8, 151.6, 140.3, 137.9, 133.9, 130.9, 129.4, 125.0, 124.7, 123.6, 117.6, 116.8, 115.2, 114.4, 110.4, 21.8, 19.9, 12.8. HRMS (ESI) for $C_{21}H_{18}F_2N_4SNa$: Calculated: 392.1118, found: 392.1354; Purity by HPLC-UV (214 nm)-ESI-MS: 98%. mp 206-207 °C.

**N-(5-Fluoro-2,4-dimethylphenyl)-4-(3-fluoro-4-(4-methyl-1H-imidazol-1-yl)phenyl)thiazol-2-amine (26).** Synthesized using N-((5-fluoro-2,4-dimethylphenyl)carbamothioyl)benzamide (**I**, 100 mg, 0.34 mmol) and 1-(5-fluoro-2,3-dimethylphenyl)thiourea (**60**, 67.40 mg, 0.34 mmol). Yield (104.80 mg, 84%). $^{1}$H NMR (600 MHz, DMSO-$d_6$) $\delta$ 9.85 (s, 1H), 8.23 (dd, $J$ = 7.5, 5.1 Hz, 1H),7.87 – 7.82 (m, 1H), 7.75 (dd, $J$ = 7.5, 5.1 Hz, 1H), 7.63 (dd, $J$ = 8.1, 1.4 Hz, 1H), 7.59 –7.42 (s, 1H), 7.30 (d, $J$ = 7.9 Hz, 1H), 7.22 – 7.16 (m, 1H), 7.07 – 7.04 (m, 1H), 2.39 (d, $J$ = 0.7 Hz, 3H), 2.19 (s, 3H), 2.15 (s, 3H). $^{13}$C NMR (150 MHz, DMSO-$d_6$) $\delta$ 162.5, 158.1, 155.6, 153.8, 151.6, 140.3, 135.7, 130.9, 128.6, 126.7, 126.4, 122.3, 122.1, 120.5, 116.8, 113.8, 110.4, 18.1, 16.7, 12.9. HRMS (ESI) for $C_{21}H_{18}F_2N_4SNa$: Calculated: 392.1118, found: 392.1905; Purity by HPLC-UV (214 nm)-ESI-MS: 96%. mp 206-207 °C.

**N-(5-Ethyl-2-fluoro-4-methylphenyl)-4-(3-fluoro-4-(4-methyl-1H-imidazol-1-yl)phenyl) thiazol-2-amine (27).** Synthesized using N-((5-fluoro-2,4-dimethylphenyl)carbamothioyl) benzamide (**I**, 100 mg, 0.34 mmol) and 1-(5-ethyl-2-fluoro-4-methylphenyl)thiourea (**61**, 72.0 mg, 0.34 mmol). Yield (192 mg, 72%). $^{1}$H NMR (600 MHz, DMSO-$d_6$) $\delta$10.29 (s, 1H), 8.30 – 8.22 (m, 1H), 8.17 (dd, $J$ = 7.5, 5.1 Hz, 1H), 7.87 (dd, $J$ = 8.1, 1.4 Hz, 1H), 7.66 – 7.62 (m, 2H), 7.42 – 7.36 (m, 2H), 7.14 (d, $J$ = 8.0 Hz, 1H), 2.51 (qd, $J$ = 8.0, 1.0 Hz, 2H), 2.37 (d, $J$ = 0.7 Hz, 3H), 2.21 (s, 3H), 1.13 (t, $J$ = 8.0 Hz, 3H). $^{13}$C NMR (150 MHz, DMSO-$d_6$) $\delta$ 162.5, 155.6, 154.9, 153.8, 151.6, 140.3, 138.6, 137.9, 135.1, 130.9, 128.6, 126.7, 124.2, 122.1, 116.2, 110.4, 20.1, 19.4, 19.1, 12.9. HRMS (ESI) for $C_{22}H_{20}F_2N_4SNa$: Calculated: 433.1275, found: 433.1183; Purity by HPLC-UV (214 nm)-ESI-MS: 97%. mp 209-210 °C.

**4-(3-Fluoro-4-(4-methyl-1H-imidazol-1-yl)phenyl)-N-(2-fluoro-4-methylphenyl)thiazol-2-amine (28).** Synthesized using N-((5-fluoro-2,4-dimethylphenyl)carbamothioyl)benzamide (**I**, 100 mg, 0.34 mmol) and 1-(2-fluoro-4-methylphenyl)thiourea (**62**, 62.50 mg, 0.34 mmol). Yield (113 mg, 88%). $^{1}$H NMR (600 MHz, DMSO-$d_6$) $\delta$ 9.50 (s, 1H), 8.16 (dd, $J$ = 7.5, 5.1 Hz, 1H), 8.01 (dd, $J$ = 8.1, 1.4 Hz, 1H), 7.79 (s, 1H), 7.68 (dd, $J$ = 7.5, 5.0 Hz, 1H), 7.42 – 7.36 (m, 2H), 7.18 – 7.07 (m, 1H), 6.89 (m, 1H), 2.37 (d, $J$ = 0.6 Hz, 3H), 2.29 (s, 3H). $^{13}$C NMR (150 MHz, DMSO-$d_6$) $\delta$ 162.5, 155.6, 154.1, 153.8, 151.6, 151.4, 140.3, 137.9, 134.3, 130.9, 126.7, 124.2, 122.7, 116.4, 115.2, 110.4, 20.3, 12.9. HRMS (ESI) for $C_{20}H_{16}F_2N_4SNa$: Calculated: 405.0898, found: 405.0762; Purity by HPLC-UV (214 nm)-ESI-MS: 99%. mp 208-209 °C.

**N-(4-Fluoro-2-methylphenyl)-4-(3-fluoro-4-(4-methyl-1H-imidazol-1-yl)phenyl)thiazol-2-amine (29).** Synthesized using N-((5-fluoro-2,4-dimethylphenyl)carbamothioyl)benzamide (**I**, 100 mg, 0.34 mmol) and 1-(4-flouro-2-methylphenyl)thiourea (**63**, 62.50 mg, 0.34 mmol). Yield (113 mg, 88%). $^{1}$H NMR (600 MHz, DMSO-$d_6$) $\delta$ 9.57 (s, 1H), 8.23 (dd, $J$ = 12.2, 2.7 Hz, 1H), 7.96 (s, 1H), 7.91 (dd, $J$ = 12.6, 5.1 Hz, 1H), 7.84 (dd, $J$ = 8.3, 1.3 Hz, 1H), 7.61 (s, 1H), 7.31 (s, 1H), 7.28 – 7.19 (m, 1H), 6.86 – 6.80 (m, 1H), 2.29 (s, 3H), 2.19 (d, $J$ = 1.1 Hz, 3H). $^{13}$C NMR (150 MHz, DMSO-$d_6$) $\delta$ 164.5, 162.1, 160.2, 155.6, 153.6, 148.0, 138.1, 136.9, 135.5, 131.9, 125.9, 124.4, 123.0, 122.7, 116.6, 114.1, 108.9, 106.6, 18.0, 13.9. HRMS (ESI) for $C_{20}H_{16}F_2N_4SNa$: Calculated: 405.0898, found: 405.1145; Purity by HPLC-UV (214 nm)-ESI-MS: 98%. mp 207-208 °C.

**4-(3-Fluoro-4-(4-methyl-1H-imidazol-1-yl)phenyl)-N-(2-fluoro-5-methylphenyl)thiazol-2-amine (30).** Synthesized using N-((5-fluoro-2,4-dimethylphenyl)carbamothioyl)benzamide (**I**, 100 mg, 0.34 mmol) and 1-(2-fluoro-5-methylphenyl)thiourea (**64**, 62.50 mg, 0.34 mmol). Yield (113 mg, 88%). $^{1}$H NMR (600 MHz, DMSO-$d_6$) $\delta$ 9.51 (s, 1H), 8.21 (dd, $J$ = 7.5, 5.1 Hz, 1H), 7.93 (dd, $J$ = 8.1, 1.4 Hz, 1H), 7.74 – 7.71 (m, 2H), 7.54 (s, 1H), 7.22 – 7.08 (m, 2H), 6.89 – 6.84 (m, 1H), 2,29 (s, 3H), 2.19 (d, $J$ = 1.5 Hz, 1H). $^{13}$C NMR (150 MHz, DMSO-$d_6$) $\delta$ 162.5, 157.1, 155.6, 153.8, 151.6, 140.3, 137.9, 134.9, 130.3, 127.1, 126.7, 123.6, 122.3, 116.2, 115.2, 110.4, 20.5, 12.9. HRMS (ESI) for $C_{20}H_{16}F_2N_4SNa$: Calculated: 405.0898, found: 405.4129; Purity by HPLC-UV (214 nm)-ESI-MS: 99%. mp 206-207 °C.

**N-(5-Fluoro-2-methylphenyl)-4-(3-fluoro-4-(4-methyl-1H-imidazol-1-yl)phenyl)thiazol-2-amine (31).** Synthesized using N-((5-fluoro-2,4-dimethylphenyl)carbamothioyl)benzamide (**I**, 100 mg, 0.34 mmol) and 1-(5-fluoro-2-methylphenyl)thiourea (**65**, 62.50 mg, 0.34 mmol). Yield (113 mg, 88%). $^{1}H$ NMR (600 MHz, DMSO-$d_6$) δ 9.57 (s, 1H), 8.23 (dd, *J* = 12.2, 2.7 Hz, 1H), 7.95 (t, *J* = 16, 1.4 Hz, 1H), 7.91 (dd, *J* = 12.6, 1.9 Hz, 1H), 7.87 – 7.80 (m, 1H), 7.69 – 7.61 (m, 1H), 7.58 (s, 1H), 7.34 (dt, *J* = 2.5, 1.5 Hz, 1H), 7.22 – 7.13 (m, 3H), 6.77 – 6.72 (m, 1H), 2.29 (s, 3H), 2.19 (d, *J* = 1.1 Hz, 3H). $^{13}C$ NMR (150 MHz, DMSO-$d_6$) δ 163.3, 161.3, 155.7, 147.4, 145.5, 137.4, 136.8, 134.8, 131.5, 130.8, 128.7, 126.7, 122.6, 121.9, 120.6, 118.2, 113.9, 22.5, 14.1. HRMS (ESI) for $C_{20}H_{16}F_2N_4SNa$: Calculated: 405.0898, found: 405.1643; Purity by HPLC-UV (214 nm)-ESI-MS: 99%. mp 205-206 °C.
